# Genotype sampling for deep-learning assisted experimental mapping of fitness landscapes

**DOI:** 10.1101/2024.01.18.576262

**Authors:** Andreas Wagner

## Abstract

**Motivation:** Experimental characterization of fitness landscapes, which map genotypes onto fitness, is important for both evolutionary biology and protein engineering. It faces a fundamental obstacle in the astronomical number of genotypes whose fitness needs to be measured for any one protein. Deep learning may help to predict the fitness of many genotypes from a smaller neural network training sample of genotypes with experimentally measured fitness. Here I use a recently published experimentally mapped fitness landscape of more than 260,000 protein genotypes to ask how such sampling is best performed.

**Results:** I show that multilayer perceptrons, recurrent neural networks (RNNs), convolutional networks, and transformers, can explain more than 90 percent of fitness variance in the data. In addition, 90 percent of this performance is reached with a training sample comprising merely ≈10^3^ sequences. Generalization to unseen test data is best when training data is sampled randomly and uniformly, or sampled to minimize the number of synonymous sequences. In contrast, sampling to maximize sequence diversity or codon usage bias reduces performance substantially. These observations hold for more than one network architecture. Simple sampling strategies may perform best when training deep learning neural networks to map fitness landscapes from experimental data.

## Introduction

A fitness or adaptive landscapes is a high-dimensional analogue to a landscape in physical space. Each genotype of an organism or biomolecule corresponds to a spatial location, and the elevation at that location corresponds to fitness. Darwinian evolution can be viewed as an exploration of such a landscape that drives evolving populations towards high fitness peaks (Wright, 1932). Characterizing the topography of a fitness landscape and identifying its highest peaks is central to both evolutionary biology and biomedical engineering.

The first experimental data on fitness landscapes became available only in the early 2000s, when experimental measurements of multiple mutations in the antibiotic resistance protein TEM-1 beta lactamase showed that few mutational paths to high antibiotic resistance are evolutionarily accessible, i.e., fitness-increasing for each mutational step (Weinreich, et al., 2006). Since then, numerous experimental studies on the topography of adaptive landscapes have been published, some based on few genotypes (Chou, et al., 2011; de Visser and Krug, 2014; Hall, et al., 2010; Mira, et al., 2015; Palmer, et al., 2015; Weinreich, et al., 2018; Yang, et al., 2019), others based on thousands (Bendixsen, et al., 2019; Diss and Lehner, 2018; Domingo, et al., 2018; Li, et al., 2016; Li and Zhang, 2018; Olson, et al., 2014; Poelwijk, et al., 2019; Pokusaeva, et al., 2019; Sarkisyan, et al., 2016). Some studies directly quantify organismal fitness conveyed by different genotypes(Chou, et al., 2011; Domingo, et al., 2018; Hall, et al., 2010; Li, et al., 2016; Li and Zhang, 2018; Palmer, et al., 2015). Many others quantify molecular traits that can serve as a proxy for fitness, such as gene expression (Li, et al., 2019), enzyme activity (Bendixsen, et al., 2019; Yang, et al., 2019), light emission by fluorescent proteins (Poelwijk, et al., 2019; Sarkisyan, et al., 2016; Zheng, et al., 2019), strength of protein-protein interactions (Diss and Lehner, 2018; Olson, et al., 2014; Wu, et al., 2016), protein-RNA binding(Melamed, et al., 2013), or protein-DNA binding (Aguilar-Rodriguez, et al., 2017).

To study the topography of adaptive landscapes is challenging. First, current theoretical models cannot predict the fitness in such landscapes from genotype alone (Das and Krug, 2022; de Visser and Krug, 2014; Kauffman and Levin, 1987; Weinreich, et al., 2018), because different base pairs interact in complex non-additive ways to determine a genotype’s fitness (Domingo, et al., 2018; Poelwijk, et al., 2019; Poelwijk, et al., 2011; Weinreich, et al., 2018; Weinreich, et al., 2013; Weinreich, et al., 2005; Yang, et al., 2019). Second, adaptive landscapes have astronomical sizes. For example, even a small gene of 100 base pairs has 4^100^=1.6×10^60^ possible genotypes. The largest landscapes mapped to date have 10^5^-10^7^ characterized genotypes (Bendixsen, et al., 2019; Chou, et al., 2011; Domingo, et al., 2018; Li, et al., 2016; Papkou, et al., 2023; Pokusaeva, et al., 2019; Sarkisyan, et al., 2016; Vaishnav, et al., 2022).

Given these challenges, it would be highly desirable to use machine learning methods to help map otherwise prohibitively large landscapes. This would involve a three-step process. First, experimentally measure the fitness of a manageable sample of genotypes from a landscape. Second, use the resulting data as training and validation data for a machine learning algorithm to predict genotype fitness. Third, test this prediction by experimentally measuring the fitness of additional genotypes as a test set. If the algorithm generalizes well to the test set, it can be used to study the topography of the entire landscape.

Machine learning in general, and deep learning in particular have proven highly successful in predicting biological phenomena (Adrion, et al., 2020; Alipanahi, et al., 2015; Alley, et al., 2019; Avsec, et al., 2021; Fernandez-de-Cossio-Diaz, et al., 2021; Flagel, et al., 2019; Govindarajan, et al., 2015; Rao, et al., 2019; Riesselman, et al., 2018; Romero, et al., 2013; Tareen, et al., 2022; Vaishnav, et al., 2022; Washburn, et al., 2019; Xue, et al., 2021; Zhou, et al., 2022). They can predict gene expression (Vaishnav, et al., 2022; Washburn, et al., 2019), protein structure and pathogenicity (Cheng, et al., 2023; Jumper, et al., 2021), protein stability (Blaabjerg, et al., 2023; Pancotti, et al., 2021), protein-nucleic acid binding (Alipanahi, et al., 2015; Avsec, et al., 2021), DNA methylation (Angermueller, et al., 2017), mutational effects on proteins and RNA (Riesselman, et al., 2018), ribosomal binding site activity (Hollerer, et al., 2020), recombination rates and selective sweeps (Adrion, et al., 2020; Flagel, et al., 2019; Xue, et al., 2021). Furthermore, they can help to classify proteins into families (Asgari and Mofrad, 2015), and to generate protein sequences with desired functions (Madani, et al., 2023).

Several studies have used machine learning to predict molecular phenotypes that can be correlated with fitness, such as enzyme activity, gene expression, and the affinity of transcription factors to DNA (Alley, et al., 2019; Li, et al., 2019; Tareen, et al., 2022; Vaishnav, et al., 2022; Wittmann, et al., 2021; Xu, et al., 2020). The application of machine learning most closely related to the present work aims at reducing the screening effort of directed evolution experiments (Li, et al., 2019; Wittmann, et al., 2021; Wu, et al., 2019). Such experiments usually improve enzymes through multiple rounds of mutagenesis and screening of variants with high “fitness”, such as a faster rate of enzymatic catalysis. The screening step in directed evolution is usually manual, labor-intensive, and thus feasible only for few (10^1^-10^2^) variants. In addition, most screened variants are inviable – have very low fitness -- and need to be excluded from future rounds, which amounts to experimental effort wasted on these variants (Li, et al., 2019; Wittmann, et al., 2021; Wu, et al., 2019). The most pertinent existing work is not based on neural networks or on shallow neural networks (Li, et al., 2019; Wittmann, et al., 2021; Wu, et al., 2019). It asks how sequence data is best-encoded, and how useful machine learning models can be to predict inviable genotypes when trained on very small samples. For example, one study (Wittmann, et al., 2021) found that a simple one-hot encoding or an encoding based on physicochemical amino acid properties performs equally well or better than sophisticated encodings pre-learned on vast data sets (Elnaggar, et al., 2021; Iuchi, et al., 2021; Rao, et al., 2021; Rives, et al., 2021).

This contribution differs from previous efforts in several ways. First, the experimental landscape data set I analyze is large, quantifies fitness in vivo, and contains data on many synonymous DNA sequences (Papkou, et al., 2023). The latter property is important, because elimination of synonymous sequences is central to some strategies to sample genotypes for experimental fitness measurements. More specifically, I analyze data on *E.coli* fitness on the antibiotic trimethoprim for almost 4^9^≈260,000 *E. coli* genotypes that differ at nine consecutive base pairs of the gene for dihydrofolate reductase (DHFR), which can convey trimethoprim resistance. (Papkou, et al., 2023).

Second and most importantly, I study how the quality of fitness predictions of several deep-learning neural network architectures, including multilayer perceptrons, recurrent neural networks, and transformers, depends on how the training data is sampled. I show that random sampling and sampling of few synonymous DNA sequences per amino acid sequences leads to the best generalization performance on test data. In contrast, sampling maximally diverse nucleotide or amino acid sequences leads to the poorest performance.

## Methods

### Data

To predict fitness for viable genotypes by (nonlinear) regression I used the 17,774 viable genotypes of the fitness data in (Papkou, et al., 2023). This experimentally measured fitness data is a logarithmically transformed *E. coli* growth rate relative to a wild-type, which has a fitness of zero. It ranges between -1.17 and +1.4. All genotypes with fitness below -0.5 are inviable (Papkou, et al., 2023). To avoid divergence of the mean absolute percentage error (mape) for fitness values around zero, I added an offset of +2 to all fitness values before training, so that they range between 0.83 and 3.4 after this transformation.

### Neural network training

I trained neural networks of all architectures with the minibatch gradient descent method, using a batch size of 128 genotypes (Bertsekas, 1996). To this end, I employed the widely used root mean square propagation (rmsprop) algorithm, as implemented in keras (tensorflow version 2.12.0, https://github.com/tensorflow/tensorflow/releases)(Chollet, 2021) I tuned hyperparameters with a hyperband tuner implemented in tensorflow (version 2.12.0, tuner parameters: factor=3, hyperband_iterations=3) (Li, et al., 2017). I used this hypertuner for 10 epochs per network, but stopped training for any one network when training showed no further improvement in performance for 5 epochs (Chollet, 2021). See Supplementary Methods for details on the network architectures and the tuned hyperparameters.

### Genotype sampling

I also restricted genotype sampling to the 17,774 viable genotypes, which encode 1,630 unique amino acid sequences. For random (uniform) sampling of genotypes, I first randomly shuffled all viable genotypes and set aside 50 percent (8887) of them as a test set, and the remainder for validation and training. I then sampled a fixed number of the remaining genotypes for training and validation. I varied this number between *S*=200 (1.1 percent of all data) and *S*=8,000 (45 percent) to explore how prediction quality depends on *S*. Because many of the resulting training/validation data sets were small, I did not use hold-out validation, but applied 4-fold cross-validation, setting aside 75 percent of the sample for training and 25 percent for validation, and repeating this procedure four times with non-overlapping validation data sets for each replicate. I computed the training and validation loss (mean squared error, mse, of predicted fitness) after each epoch as an average across the four training runs.

For each training sample I trained each network with the rmsprop algorithm for a maximum of 100 epochs with batch sizes of 128 samples. I stopped the training early when the training loss (mse) did not decrease for five consecutive epochs. I trained each network in three independent replicates to estimate how much fitness predictions vary across such replicates. I chose independent test and training/validation data sets for each value of *S* and for each replicate. I used the same procedure also for the non-random sampling procedures described in the text (Supplementary Methods).

## Results

### Recurrent neural networks are best at predicting the fitness of viable genotypes

Just like for other proteins (Li, et al., 2019; Wittmann, et al., 2021; Wu, et al., 2019), only a small minority of the genotypes (17,774, 6.8 percent) in the DHFR trimethoprim resistance landscape is viable (Papkou, et al., 2023). I study the ability of 6 neural network architectures to distinguish viable from inviable genotypes (Supplementary Results 1) and to predict the fitness of these viable genotypes by (nonlinear) regression.

As one of two base-line reference models to predict fitness, I use a random predictor. This predictor uses fitness values that are randomly shuffled among genotypes. It performs poorly, predicting less than 0.01 percent of the variation in fitness (Table 1). My second base-line reference model is linear regression, which already performs vastly better than random prediction, halving the mean errors (mean absolute percentage error, mape=15.65%, mean absolute error, mae=0.33), and increasing the correlation coefficients to *r*=0.66 and *R^2^*=0.41. In other words, linear regression can explain 41 percent of the variation in the data.

**Table 1:**
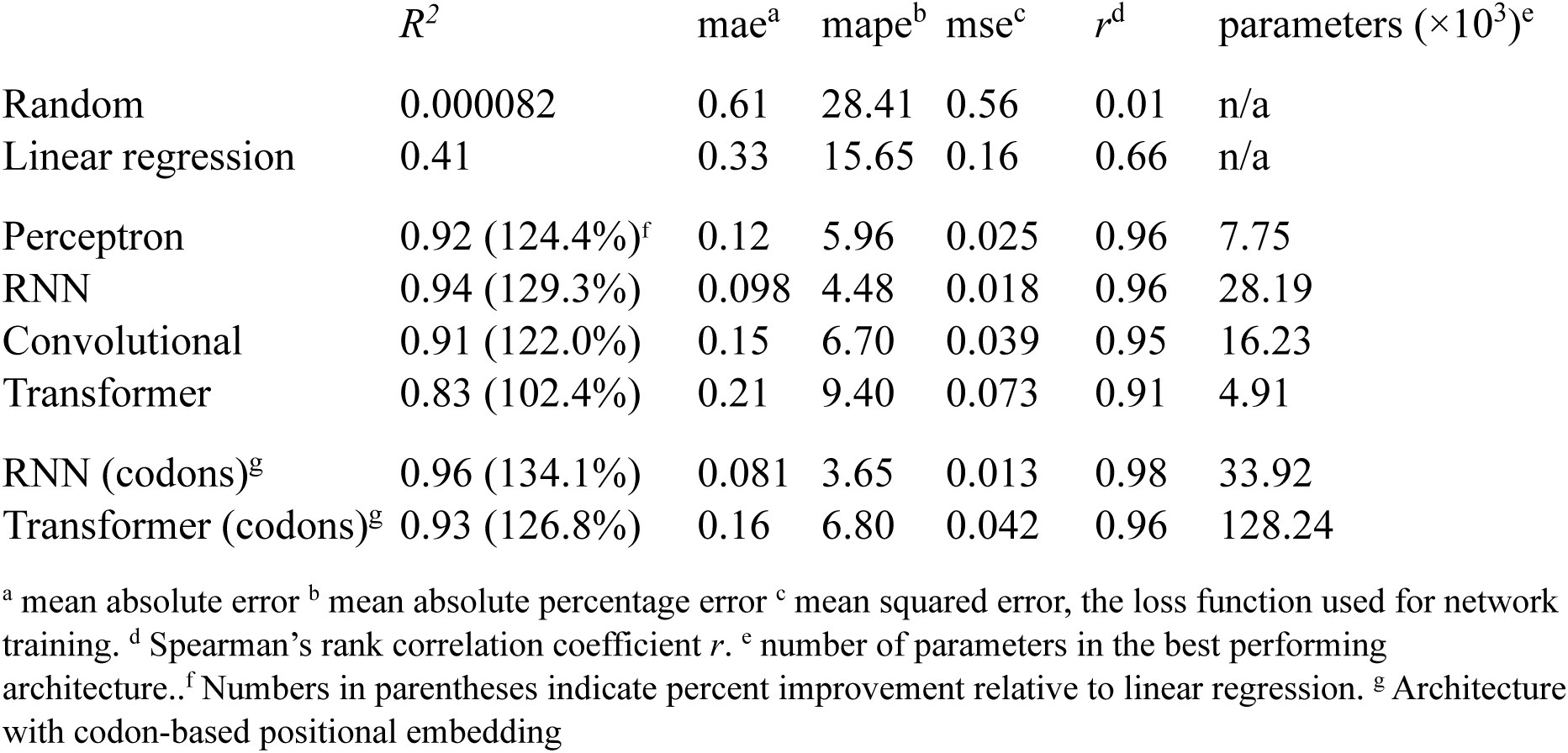
Performance of deep learning network architectures on regression of viable genotypes.

The first neural network architecture I study is the multilayer perceptron (Gurney, 1997; LeCun, et al., 2015; Rosenblatt, 1958), in which I tuned the number of layers, the number of neurons per layer, weight regularization, layer dropout, and the learning rate (Supplementary Methods). It already leads to a massive further improvement over linear regression. For example, it reduces the mape by 61.9 percent to 5.96%, and increases *R^2^*by 124.4% to *R^2^*=0.92. (See Table 1 for the other performance measurements) That is, even this simple architecture can explain 92% of the variation in the data.

The second architecture is a bidirectional recurrent neural network (RNN) (Hochreiter and Schmidhuber, 1997), in which I tuned the number of bidirectional layers, the number of neurons in each layer, weight regularization, recurrent dropout, and the learning rate. This network performed slightly better than the perceptron, with *R^2^*=0.94 (129.3% improvement over linear regression) and a mape of 4.48 percent.

The third architecture is a one-dimensional convolutional network (LeCun, et al., 2015), in which I tuned the number of convolutional layers, the number of dense layers that followed them, the number of neurons in these layers, their weight regularization, and the learning rate. It performed slightly less well (R^2^=0.91, 122% improvement over linear regression) than the preceding architectures.

The input to the three architectures I discussed thus far was a flattened one-hot encoded 9×4=36-dimensional representation of a DNA genotype. In contrast, the next architecture is a transformer (Vaswani, et al., 2017), for which I first positionally embedded individual DNA sequences in a low-dimensional embedding space (Chollet, 2021, p 347), which ensures that the embedding of each sequence also contains information about the position of each nucleotide in the sequence. The optimal embedding is learned during neural network training.

I deliberately chose such end-to-end learning of word embedding, because it performs at a par with highly complex pre-trained embeddings, may require lower embedding dimensions, and does not depend on other bioinformatic resources (Alley, et al., 2019; Asgari and Mofrad, 2015; ElAbd, et al., 2020; Elnaggar, et al., 2021; Iuchi, et al., 2021; Raimondi, et al., 2019; Rao, et al., 2021; Rives, et al., 2021).

In this transformer architecture, I tuned the number of embedding dimensions, the number of attention heads per transformer module, the size of each attention head, the number of neurons in each dense layer of a module, the number of stacked transformer modules, the dropout rate, and the learning rate (Supplementary Methods). Despite such extensive hypertuning, the transformer too performed less well than the RNN (*R^2^*=0.83, Table 1).

Feature engineering, i.e., choosing an appropriate representation of input data, can be crucial to improve network performance (Chollet, 2021). For two further neutral networks, I chose a simple and general form of feature engineering with the advantage that it would apply to all protein-coding genes and is not specific to DHFR or a specific protein class. Specifically, I subdivided the 9 nucleotide input sequence into 3 integer-encoded codons and positionally embedded these codons into a space whose dimensionality I varied during hypertuning (Supplementary Methods). These codons became the input to a bidirectional RNN whose hyperparameters I also tuned (Supplementary Methods).

Feature engineering improves the performance of the transformer by a further 12.0% to *R^2^*=0.93), as well as that of the RNN by 2.1% to *R^2^*=0.96 (Table 1). Overall, the bidirectional RNN network (Figure S13) with a codon-based embedding performs best, explaining 96 percent of the variation in fitness (mae= 0.081, mape=3.65).

With 33,921 parameters the bidirectional RNN is more complex than the simpler and almost equally well-performing multilayer perceptron (7,745 parameters, Table 1, Supplementary Results 2). The best-performing transformer requires many more parameters (128,241) despite its poorer performance. I focused my subsequent analyses on the best-performing RNN, but also compared their outcome with the best-performing perceptron and transformer, because of their widely varying complexity, to find out how strongly the influence of genotype sampling on prediction performance depends on the architecture. During training, all three types of networks converge rapidly (within 10 epochs) to their optimal performance (Figure S1). They show no signs of overfitting thereafter (Figure S1), suggesting that even better architectures exist.

### A small sample of training data can suffice to predict fitness with high accuracy

Because measuring fitness experimentally is laborious, the sample of genotypes measured sample should be as small as possible. This is especially important wherever high-throughput fitness measurements are infeasible (Nikolados, et al., 2022; Wittmann, et al., 2021). To find out whether accurate fitness prediction is even possible from a small sample, I first studied how the prediction quality of the best-performing RNN varies with the sample size *S* that is used for training and validation. Specifically, I varied this sample size between *S*=200 and *S*=8,000 randomly chosen genotypes (1.1% - 45.0% of all viable genotypes). For any one value of *S*, I subdivided all 17,774 viable genotypes into a test set that comprised 50% of the data (8,886 sequences), and a set for training and validation that comprised *S* sequences, using 4-fold cross-validation during training. Subsequently, I tested the model thus trained on the test set.

Figure 1 shows the coefficient of determination *R^2^* of predicted fitness on the test set as a function of sample size for the best-performing RNN (Figure S13), and for random (uniform) sampling of the training data. *R^2^* increases rapidly with sample size *S*, and reaches 90% of the *R^2^* obtained for the maximal sample size after training on only 7.8% (1,400) of genotypes.

**Figure 1:**
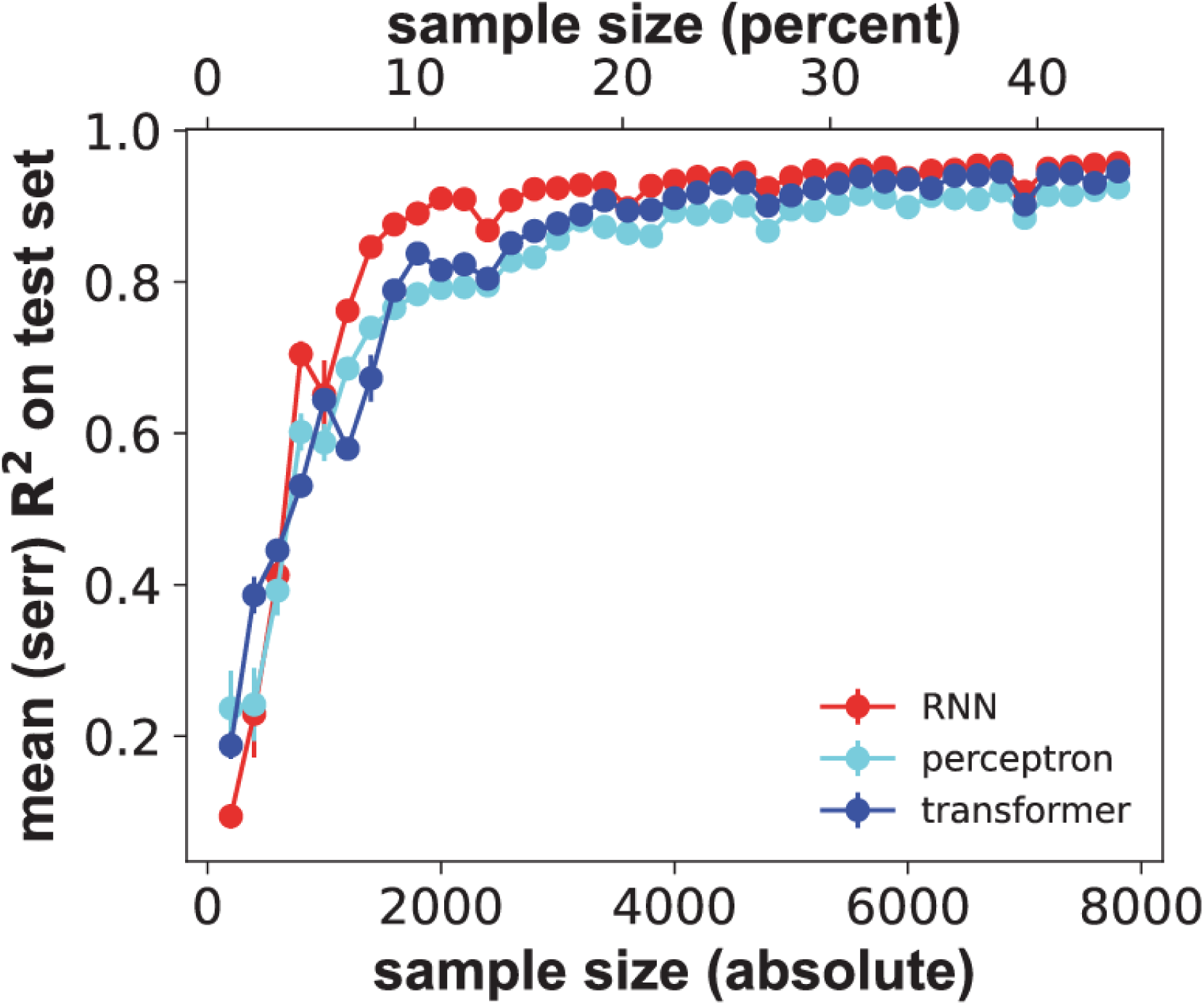
Predictive performance increases rapidly with sample size for three different neural network architectures. horizontal axis: sample size *S* of a random genotype sample used for training and validation through 4-fold cross-validation, both in absolute numbers of genotypes (bottom) and as a percentage of all viable genotypes (top). Vertical axis: performance of the three major network architectures (color legend) on a test-set comprising 50 percent (8887) of viable genotypes, as quantified by the coefficient of determination *R^2^*, between measured and predicted fitness. Whiskers indicate one standard error of the mean based on three replicate trainings for each network and sample size.

Notably, the sample sizes needed to reach a value of *R^2^*within 90% of that for the largest training set are similarly small for the multilayer perceptron and for the transformer (1,600 genotypes, 9.0 percent of all genotypes for both, Figure 1). Other measures of performance also reach close to peak performance with small samples (Figure S2). In sum, accurate fitness prediction is possible with small training sets, independently of network architecture.

### Sampling strategies that reduce the number of synonymous sequences alter performance only slightly

Random (uniform) samples of DNA sequences for fitness measurements have a key disadvantage. Because of the redundancy of the genetic code, many sampled DNA sequences will be synonymous, encoding the same amino acid sequences. Because fitness differences between synonymous sequences are usually much smaller than between non-synonymous sequences, laborious fitness measurements for synonymous sequences can waste valuable experimental resources (Bailey, et al., 2021; Cuevas, et al., 2011; McDonald and Kreitman, 1991).

These observations raise the question how much predictive power a deep learning network loses when sampling few or no synonymous sequences for each amino acid sequence. To answer this question, I first implemented a sampling procedure (‘one syn.’) that aims to create training/validation data sets in which every amino acid is only represented by a single nucleotide sequence, thus avoiding synonymous sequences altogether. Because all viable 17,774 DNA sequences encode only 1,630 amino acid sequences, synonymous sequences can only be avoided entirely for small samples (Supplementary Methods). However, the procedure creates a mean number of nucleotide sequences per amino acid sequences that is much smaller than for random samples (e.g., *S*=1,400: ‘one syn.’ creates 1.06±0.004 [mean±1 standard error] synonymous sequences per amino acid sequences; random: 2.04±0.03 synonymous sequences.

I hypothesized that this sampling method leads to better predictions than random sampling, because it samples the most informative nucleotide sequences, i.e., those that encode different proteins. However, this is not the case (Figure 2a and 2b). Here and below, I compare sampling performance mostly at *S*=1,400, because this is where the RNN first reaches 90% of its peak performance, i.e., its performance for the largest training sample. This is also where different architectures show the clearest performance differences (Figure 1). At this sample size, the mape of the RNN increases by 10 percent (to 8.25±0.44) for ‘one syn.’ sampling relative to random sampling (7.49±0.16, Figure 2c), and the mean *R^2^* decreases by 5.9 percent (to 0.80 from 0.85, Figure 2d). Likewise, this sampling method does not lead to a consistent performance improvement for the other network architectures, although the performance differences are modest (Figure S3 and S4; multilayer perceptron: mape=9.8±0.24 and 9.53±0.09; *R^2^*=0.74 to 0.71; transformer: mape=11.1±0.92 and 11.9±0.29; *R^2^*=0.67 and 0.63, each pair of numbers for random sampling and ‘one syn.’ respectively).

**Figure 2:**
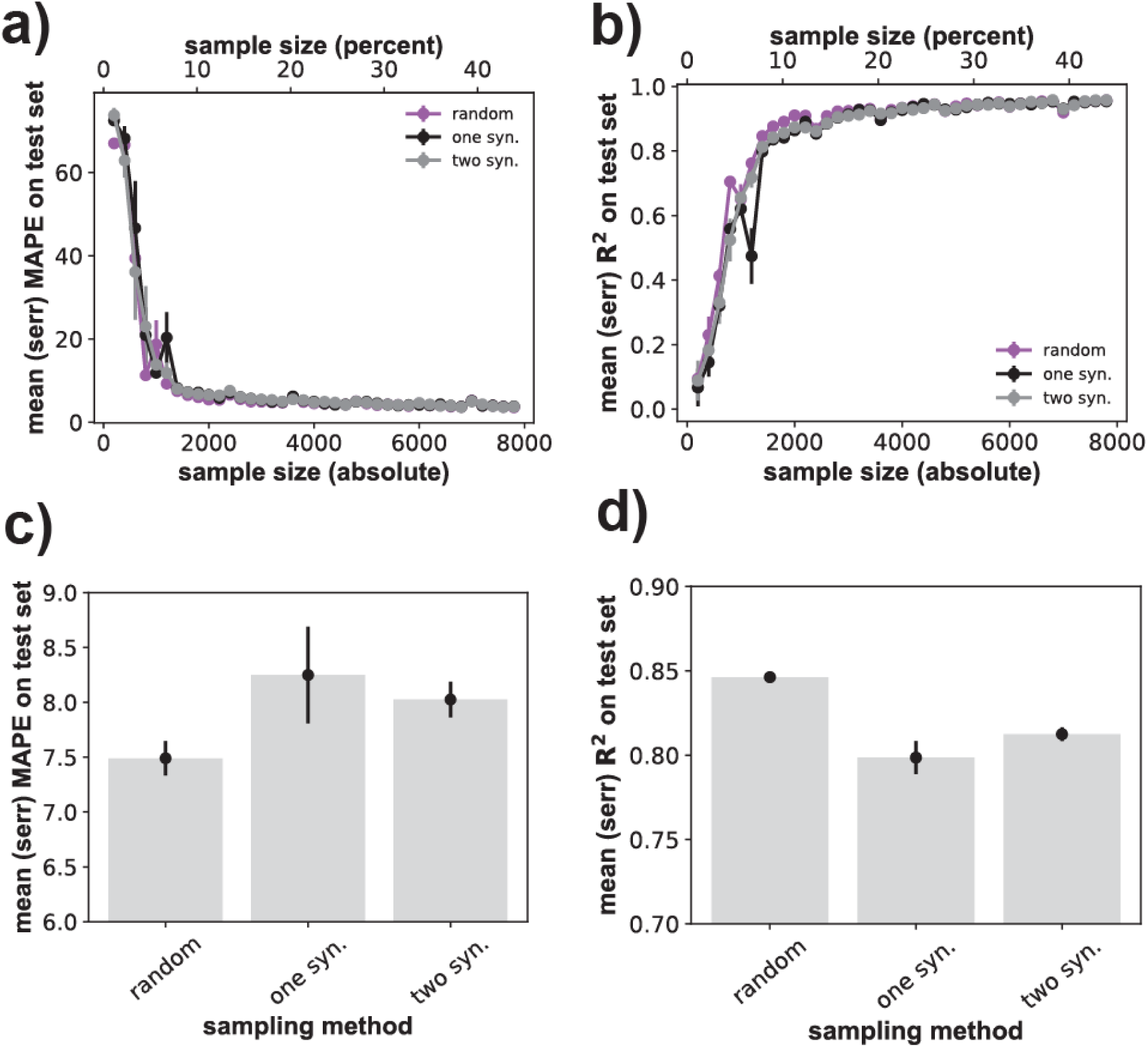
Sampling one or two synonymous sequences moderately degrades RNN prediction quality. **a)** horizontal axis: size *S* of the genotype sample used for training and validation through 4-fold cross-validation, both in absolute numbers of genotypes (bottom) and as a percentage of all viable genotypes (top). Vertical axis: prediction quality of the (best-performing) RNN architecture, as quantified by the mape of fitness prediction as a function of sample size *S*. The *S* genotypes are either sampled randomly and uniformly (‘random’), or such that only one synonymous (‘one syn.’) or two synonymous (‘two syn.’) nucleotide sequences are sampled per amino acid sequence (Supplementary Methods). Whiskers indicate one standard error of the mean, based on three replicate trainings for each network and sample size. **b)** like a), but prediction quality is quantified through the coefficient of determination *R^2^*. **c)** Dot-whisker plot indicating the means (height of bars) and standard errors (whiskers) of the mape at a fixed sample size of *S*=1,400 genotypes for the three sampling methods shown on the horizontal axis. **d)** like c), but for *R^2^* instead of the mape.

A next, less extreme sampling method (‘two syn.’) aims to create samples where every amino acid sequence is encoded by two randomly chosen nucleotide sequence. The exception is amino acid sequences that are presented by only a single encoding nucleotide sequence in the data, and large samples, where the smallest number of nucleotide sequences beyond two is sampled per amino acid sequence (Supplementary Methods). The rationale for this procedure is that it may be necessary to capture at least some of the diversity of synonymous sequences to predict fitness most accurately. (Ideally, one would sample synonymous sequences that differ in fitness, but this is not possible, because genotype fitness is unknown at the time of sampling.)

This method performs similar to ‘one syn.’ sampling (Figure 2). Specifically, at *S*=1,400 the RNN’s mape is 8.03±0.16, as compared to 8.25±0.44 for ‘one syn.’, and its *R^2^*equals 0.81 (one syn.: 0.80). The method also leads to similar performance for the other two network architectures (Figure S3 and S4; multilayer perceptron: mape=9.53±0.09 and 10.4±0.18; *R^2^*=0.71 and 0.67; transformer: mape=11.9±0.29 and 10.5±0.43; *R^2^*=0.63 and 0.69, each pair of numbers for ‘one syn.’ and ‘two syn.’, respectively).

In sum, independent of the neural network architecture, genotype sampling of few synonymous sequences does not dramatically alter performance relative to random sampling. Other methods for codon compression (Pines, et al., 2015), i.e., reducing synonymous sampling, are discussed in Supplementary Results 3.

### Increasing sampled sequence diversity reduces predictive performance substantially

In a random (uniform) sample of DNA nucleotide sequences, some sequences may be very similar to one another. Such sequences tend to encode amino acid sequences that are identical or at least physicochemically similar, and may thus have similar fitness (Freeland and Hurst, 1998). It may be best to avoid such highly similar sequences during neural network training, and instead sample more diverse sequence to facilitate generalization to a test data set.

I tested this hypothesis with two complementary sequence sampling procedures. The first aims to maximize nucleotide sequence diversity in a training/validation data sample (Supplementary Methods). Remarkably, this procedure performs substantially worse than random sampling (Figure 3). At a sample size of *S*=1,400 sequences, the mape of the RNN increases by 140.6 percent to 18.02±2.2 (Figure 3c), relative to random sampling (7.49±0.16). The mean *R^2^* decreases by 66.9 percent (from 0.85 to 0.29, Figure 3d). This sampling method also degrades the performance of the other network architectures to a similar extent (Figure S8 and S9).

**Figure 3:**
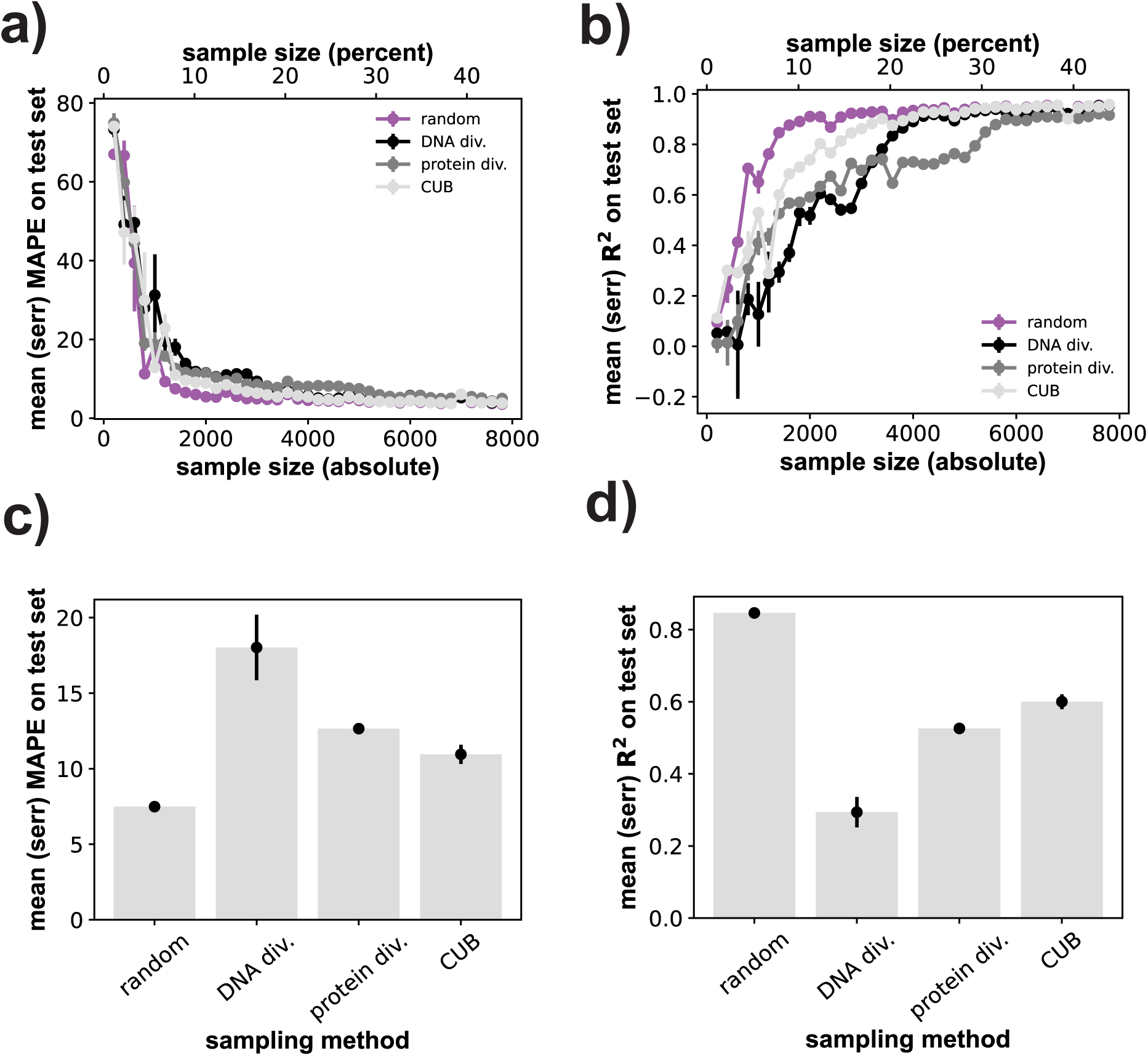
Sampling for sequence diversity or codon usage bias substantially reduces RNN prediction quality. **a)** horizontal axis: size *S* of the genotype sample used for training and validation through 4-fold cross-validation, both in absolute numbers of genotypes (bottom) and as a percentage of all viable genotypes (top). Vertical axis: prediction quality of the (best-performing) RNN architecture, as quantified by the mape of fitness prediction as a function of sample size *S*. The *S* genotypes are either sampled randomly and uniformly (‘random’), to achieve maximal DNA sequence diversity (‘DNA div.’), maximal amino acid sequence diversity (‘protein dev.’), or maximal codon usage bias (‘CUB’, Supplementary Methods). Whiskers indicate one standard error based on three replicate trainings for each network and sample size. **b)** like a), but prediction quality is quantified through the coefficient of determination *R^2^*. **c)** Dot-whisker plot indicating the means (height of bars) and standard errors (whiskers) of the mape at a fixed sample size of *S*=1,400 genotypes for the three sampling methods shown on the horizontal axis. **d)** like c), but for *R^2^*instead of the mape.

My second procedure aims to sample a set of amino acid sequences whose constituent sequences are physicochemically maximally diverse (Supplementary Methods), using a high-dimensional representation of each amino acid (Georgiev, 2009) that outperforms others in similar machine learning tasks (Wittmann, et al., 2021). This method too substantially degrades prediction quality relative to random sampling. For the RNN at a sample size of *S*=1,400, it increases the mape by 70 percent from 7.49±0.16 to 12.7±0.19 (Figure 3c). It decreases the *R^2^* by 37.6 percent from 0.85 to 0.53 (Figure 3d). Performance also declines to a similar extent for the other two architectures (Figures S8 and S9).

### Sampling sequences with high codon usage bias

I next studied a sampling procedure that preferentially samples nucleotide sequences with high codon usage bias (Supplementary Methods). Such sequences often encode proteins that are highly expressed and are thus more easily analyzed and preferred for experimental analysis (Hershberg and Petrov, 2008; Ikemura, 1985; Iriarte, et al., 2021; Komar, 2016). This procedure degrades prediction quality relative to random sampling, but more modestly than diversity-maximizing sampling. Specifically, for the RNN at a sample size of *S*=1,400, it increases the mape by 46.9 percent from 7.49±0.16 to 11.0±0.63 (Figure 3c), and decreases the *R^2^* by 29.4 percent from 0.85 to 0.6 (Figure 3d, S8, and S9; perceptron: mape increases by 26.5 percent from 9.8±0.24 to 12.4±0.77; *R^2^* decreases by 27.0 percent from 0.74 to 0.54; transformer: mape increases by 15.3 percent from 11.1±0.92 to 12.8±0.28; *R^2^* decreases by 20.9 percent from 0.67 to 0.53, all numbers for *S*=1,400). The performance differences between the sampling methods I studied also persist at much larger sample sizes, albeit at much smaller absolute performance differences (Figure S10).

## Discussion

Among the neural network architectures I studied, RNNs with a codon-based positional embedding of DNA sequences best generalized to a test data set, with a mean absolute fitness prediction error of 3.65 percent and R^2^=0.96 (Table 1, Figure S13). Remarkably, 90% of this peak performance can be reached with a training sample of merely 1,400-1,600 viable sequences (<10% of all viable sequences). This is consistent with previous observations of successful phenotype prediction from small training samples of 10^1^-10^3^ genotypes for other machine learning methods (Nikolados, et al., 2022; Wittmann, et al., 2021). This and the observations I discuss below hold not just for the RNN, but also for perceptrons and transformers. They are thus a property of the landscape and the sampling regime rather than of a specific neural network architecture.

Sampling few synonymous DNA sequences per amino acid sequence leads to the best generalization after random sampling. This observation is easily explained by the weak fitness effects of synonymous mutations (Bailey, et al., 2021; Cuevas, et al., 2011; McDonald and Kreitman, 1991), which means that synonymous DNA sequences account for less fitness variation than non-synonymous sequences.

In contrast to random sampling, sampling genotypes for highly diverse DNA sequences or highly physicochemically diverse amino acid sequences substantially degrades generalization ability. Sampling for diversity disfavors sequences within local neighborhoods. Random sampling from a small sequence space like the one I study here will cause at least some sampled sequences to lie close to each other. My observations show that such highly local sampling is important for accurate fitness predictions.

This assertion is consistent with theoretical work that examined the ability of quadratic regression models to predict the fitness of RNA molecules, as determined by a biophysically motivated algorithm for RNA secondary structure folding (du Plessis, et al., 2016). The authors studied an enormous sequence space centered on a 217 nucleotide long biological RNA sequence with variant sequences differing at up to 100 nucleotides from this center.

They showed that for a given sample size (≈10^4^) of RNA sequences used to train a regression model, prediction quality decreased as the training sequences were allowed to differ in more and more nucleotides from the reference (du Plessis, et al., 2016). The reason is that accurate fitness prediction requires sufficiently dense local sampling of sequence space, and as the radius of the sampled region increases, such sampling becomes infeasible.

Another pertinent study examined the role of sampled sequence diversity to predict the translation efficiency of a bacterial fluorescent reporter gene with deep learning models (Nikolados, et al., 2022). The 200,000 96nt sequences in this study were organized around 56 seed sequences that are distant from each other in the large space of 4^96^≈6×10^57^ DNA sequences of this length (Cambray, et al., 2018). Each of these seed sequences was mutagenized to create a local ‘cloud’ of ≈4000 sequences around the seed whose translation efficiency was measured. The study showed that training a deep learning neural network only on the sequences near one seed yields poor generalization for test data derived from sequences far from the seed. Performance substantially improved when data from an increasing number of seeds was used in training, even if the total number of sequences in the training data was held constant (Nikolados, et al., 2022).

The apparent discrepancy to my observation that diverse sequence samples lead to poor generalization can be easily explained by the smaller region of sequence space I sample. In much larger sequence spaces, sampling diverse sequences may become essential to ensure generalization to unseen sequences. The optimal balance between ‘global’ sampling of distant sequences and ‘local’ sampling around these distant sequences remains an important task for future work.

In addition to sampling diverse genotypes, sampling genotypes for favorable codon usage also substantially degrades generalization ability. One candidate explanation is that such sampling may reduce the variation of fitness in a sample, because it reduces expression variation as a contributor to fitness variation. However, this is not the case, because genotype samples with high codon usage do not vary less in fitness than random samples. (e.g., fitness standard deviation sd in three samples of *S*=1,400 genotypes: sd=0.56±0.002 when sampling for high codon usage bias, and sd=0.53±0.007 for a random sample.) To explain why training samples with high codon usage bias leads to low generalization ability remains another task for future work.

The small sequence space of the experimental fitness landscape I study is one main limitation of my work. Another is that I study only one landscape, because it is the only one currently available with not just many genotypes but many synonymous genotypes. Other landscapes may require different kinds of sampling regimes. For example, a landscape of mRNAs translational efficiency is affected by multiple and heterogenous factors, including mRNA secondary structure and hydrophobicity of the encoded peptide (Cambray, et al., 2018). Such a landscape may thus require more diverse sampling than the landscape of an enzyme’s catalytic activity. Until many and diverse landscapes have been studied, simple sampling regimes like random sampling or codon compression sampling will be the best starting points to train deep learning neural networks on experimentally mapped fitness landscapes.

## Acknowledgments

I would like to thank Dr. Andrei Papkou for helpful comments and discussions.

## Data Availability

The fitness landscape data analyzed here is publicly available as described previously (Papkou, et al., 2023). All code used to analyze this landscape will be made publicly available upon acceptance of this manuscript at https://github.com/andreas-wagner-uzh/fitness_landscape_sampling.

## Supplementary Methods

### Two major tasks for deep learning networks

I trained deep learning neural networks on two substantially different tasks. The first is the binary classification task to distinguish viable from non-viable genotypes, for which I used the entire data set of 261,332 genotypes and their associated fitness values (Papkou, et al., 2023). I randomly shuffled all genotypes in this data set, of which I used 50 percent as a training set, 25 percent as a (held-out) validation set, and 25 percent as a test set. During training, I used binary cross entropy as a loss function to quantify training progress and network performance (Cover and Thomas, 2006; Ramos, et al., 2018). I report the sensitivity and specificity of a classifier on the test as a measure of its ability to generalize to unseen data, because they are easy to interpret and widely used (Yerushalmy, 1947). Sensitivity refers to the fraction of genotypes correctly classified as viable among all viable genotypes. It can be mathematically expressed as TP/(TP+FN), where TP (true positives) is the number of genotypes that are correctly classified as viable, and FN (false negatives) is the number of genotypes that are incorrectly classified as inviable even though they are viable. A highly sensitive classifier has a low false negative rate, i.e., it rarely calls a genotype inviable even though it is viable. Conversely, specificity refers to the number of genotypes correctly classified as inviable. It can be calculated as TN/(TN+FP), where TN (true negatives) is the number of genotypes correctly classified as inviable, and FP (false positives) is the number of genotypes incorrectly classified as viable even though they are inviable. A highly specific classifier has a low false positive rate, i.e., it rarely calls a genotype inviable even though it is viable.

The second task was the (nonlinear) regression task to predict fitness for viable genotypes. For this task, I used only the 17,774 viable genotypes (6.8% of the whole data set), and split them after random shuffling again into a training set (50% of all genotypes), a validation set (25%), and a test set (25%). During neural networks training, I used the mean squared error (mse) between the actual and predicted fitness as a loss or objective function to be minimized, because it is differentiable around zero. However, I report the mean absolute percentage error (mape) as a measure of network performance, because it is easier to interpret. One limitation of the mape is that it diverges for values of fitness close to zero. The experimentally measured fitness data is a logarithmically transformed growth rate relative to a wild-type, which has a fitness of zero. It ranges between -1.17 and +1.4, and all genotypes with fitness below -0.5 are inviable (Papkou, et al., 2023). To avoid divergence of the mape, I added an offset of +2 to all fitness values before training, so that they range between 0.83 and 3.4 after this transformation. Another limitation of the mape is that it is sensitive to the value of this offset – it approach zero as this offset approaches infinity. I thus also report the mean absolute error (mae) as a measure of network performance in regression, which does not have this limitation. In addition, I report the squared Pearson correlation coefficient *R^2^* between actual and predicted fitness.

### Linear regression benchmarks

To compute a benchmark of genotype classification based on linear regression, I applied linear regression to a one-hot encoded flattened representation of all 261,332 genotypes to compute a predicted fitness for each genotype. I then classified a genotype as predicted to be viable (inviable) if its predicted fitness fell above (below) the viability threshold of 1.5. I compared this classification to that of the experimentally determined fitness to compute TP, TN, FN, and FP, and thus to determine the sensitivity and specificity of a classifier based on linear regression. To compute a benchmark of fitness prediction based on linear regression, I applied linear regression to a one-hot encoded flattened representation of the 17,774 viable genotypes.

### Neural network training

To train all neural networks I used minibatch gradient descent training with a batch size of 128 genotypes (Bertsekas, 1996). To this end, I employed the widely used root mean square propagation (rmsprop) algorithm to implement gradient descent training. This algorithm adjusts the step size by which network parameters are changed during training by a value that is the inverse of the square root of a moving and decaying average of past gradient values.

This procedure ensures that step size is large if recently encountered gradient values are small and vice versa. Specifically, I used the algorithm as implemented in keras (tensorflow version 2.12.0, https://github.com/tensorflow/tensorflow/releases)(Chollet, 2021)

### Neural network architectures

I trained neural networks of several architectures on both the classification and regression tasks, using hyperparameter tuning to identify features such as the number of layers to identify well-performing networks. To this end I used a hyperband tuner implemented in tensorflow (version 2.12.0), which reduces tuning time by training networks with different hyperparameters for a small number of epochs before comparing their performance (tuner parameters: factor=3, hyperband_iterations=3)(Li, et al., 2017). Specifically, I used this hypertuner for 10 epochs per network, but stopped training for any one network when training showed no further improvement in performance for 5 epochs (Chollet, 2021)

#### Stack of densely connected layers (multilayer perceptron)

The input to this network was a flattened one-hot encoded nine nucleotide DNA genotype, i.e., a binary vector of dimension 9×4=36. Each layer had *N*= 8, 16, 32, or 64 densely (fully) connected internal neurons (units) whose number I tuned. I fed the input into a single dense layer of *N* units. The output became the input of between *M*=0 and *M*=3 further modules, where each module comprised a pair of dense layers with *N* units each. I normalized the output of each two-layer module to a mean of zero by applying layer normalization, which can help to reduce training time (Xu, et al., 2019). Each layer used the rectified linear unit (relu) as an activation function (Fukushima, 1975).

For each dense layer described thus far, I regularized the layer weights, which helps to avoid overfitting by adding the product of a penalty term proportional to the sum of the squares of all layer weights to the loss function. I applied this (L2)-regularization to the kernel weights of each layer and explored penalty terms of 0, 10^-4^, and 10^-3^ during tuning. Also, for each layer, I used layer dropout, i.e., I set a fraction of input units to each but the first layer of the network to zero during tuning, which also helps to avoid overfitting. I tuned the fraction of such dropout units, exploring values of 0.0, 0.1 or 0.2. In addition, I added a residual connection between consecutive two-layer stacks, which can help reduce training errors in deep neural networks (He, et al., 2016).

I fed the output of the last layer of this architecture into a dense output layer with a single unit. This unit had a sigmoid activation function for the classification task. I classified each genotype as viable (1) or inviable (0) if the output of this function was greater or equal to 0.5 or less than 0.5, respectively. For the regression task, the single output unit had no activation function (linear activation). In sum, the networks I tuned had between 2 and 8 (=1+2*M*+1, where 0≤*M*≤3) densely connected layers.

Finally, I also tuned the learning rate of the gradient descent algorithm, exploring learning rates of 10^-4^, 10^-3^, and 10^-2^. Overall, I thus tuned 5 hyperparameters, the number *N* of units per layer, the number *M* of two-layer stacks, the regularization penalty, the dropout rate, and the learning rate. I use the term multilayer perceptron for this architecture, because it is widely employed, even though the original perceptron did not use the relu activation function (Gurney, 1997; LeCun, et al., 2015; Rosenblatt, 1958).

#### Recurrent neural networks (RNNs)

This type of architecture uses bidirectional long-short term memory recurrent neural networks (Hochreiter and Schmidhuber, 1997). I chose bidirectional and not unidirectional recurrent networks, because the context of DNA nucleotides both before and after a given nucleotide may affect fitness. The input to these networks was a one-hot encoded representation of the DNA genotype, i.e., a 9×4=36-dimensional binary vector. I fed this input into a stack of *M*=3-5 bidirectional RNNs, each with *N*=8, 16, 32, or 48 units. I tuned both *M* and *N* during hypertuning, such that all *M* RNNs had the same number *N* of units.

For each RNN, I applied kernel regularization and recurrent regularization (Zaręba, et al., 2015). I tuned the regularization penalty of both, exploring values of 0, 10^-4^, and 10^-3^. Also for each RNN, I used recurrent dropout (applied to the recurrent state), exploring dropout rates of 0, 0.1, and 0.2 (Baldi and Sadowski, 2013). For each layer after the first bidirectional layer, I applied batch normalization (Ioffe and Szegedy, 2015). In addition, for all but the first and last layers, I applied residual connections between the output of the previous layer and the output of the current layer. I then fed the output of the final recurrent layer into a dense layer of one unit for which I used again a sigmoid activation function in the classification task, and no activation function in the regression task. Finally, tuned the learning rate of the gradient descent training algorithm, exploring values of 10^-4^, 10^-3^, and 10^-2^.

In sum, I tuned 8 hyperparameters, (1) the number of RNNs, (2) the number of units in each RNN, (3) dropout within each RNN, (4) regularization penalties for each RNN, as well as (5) the learning rate.

##### Transformer

The architecture I describe below is a standard transformer architecture adapted to the task at hand (Vaswani, et al., 2017). I first positionally embedded individual DNA sequences in a lower dimensional embedding space, as described in (Chollet, 2021, p 347), which ensures that the embedding of each sequence also contains information about the position of each nucleotide in the sequence. To this end, I first integer-encoded each DNA sequence, such that each nucleotide is represented by an integer (A: 1, C: 2, G: 3, T: 4). The integer-encoded DNA sequences then becomes the input to a positional embedding layer (Chollet, 2021) whose dimension *D* I tuned, exploring values of *D*=2, 3, 4, and 8. The optimal embedding is learned during neural network training.

I then passed this data to a multi-head attention layer with identical key, query, and value pairs, which implements self-attention (Vaswani, et al., 2017). I applied residual connection by adding the output of the embedding layer to the output of the multi-head attention layer (He, et al., 2016), and applied layer normalization to the resulting data (Xu, et al., 2019). This normalized data then became the input to two consecutive dense layers. I applied residual connection once again by adding the input to the first dense layer to the output of the second dense layer, and then used again layer normalization.

Collectively, the multi-head attention layer together with the two dense layers, the residual connection, and the normalization step constitute one module of the transformer. I then stacked 1, 2, 4, or 6 such modules during tuning. In addition, for each module I tuned the number of attention heads (2, 4, 6 or 8), the key dimension, i.e., the size of each attention head (2, 3, 4, and 8), as well as the number of units in each dense layer, (4, 8, 16). Moreover, I tuned whether or not to apply dropout at a fixed rate of 0.1 to the output of the last transformer module. I flattened the resulting output and fed it into a dense layer with a single output unit and a sigmoid activation function (for classification) or no activation (for regression). I trained the resulting network at hypertuned learning rates of 10^-4^, 10^-3^, or 10^-2^.

##### One-dimensional convolutional neural network

This architecture uses as input flattened one-hot encoded genotype data, which leads to a 9×4=36-dimensional input vector for each genotype. Because each nucleotide in this representation corresponds to four consecutive entries of the input vector, the first layer used a stride of four. In addition, it used an eight-dimensional convolutional kernel, and 16 filters, which allows the layer to extract all 16 possible translation-invariant dinucleotide features that could occur in the genotype data.

I followed this layer with M=1-6 further convolutional layers, each with a stride of one and a kernel size of two, reasoning that such layers could extract ever more complex translation-invariant features if such features exist. I endowed these layers with 24, 32, 48, 64, 96, or 128 filters. That is, if I added only one such layer, it had 24 filters, if I added two such layers, the first had 24 and the second 32 filters, and so on. I did not use any pooling (Weng, et al., 1993) to reduce data dimensionality between layers, because the genotypes I studied were very short and did not require such subsampling. All convolutional layers used a rectified linear unit (relu) activation function.

I flattened the output of the last convolutional layer, and fed it into a stack of 0, 1, or 2 dense layers with 8, 16, or 32 units. I applied L2 kernel regularization to each dense layers, with a penalty that I tuned to 0, 10^-4^, and 10^-3^. I fed the output of the last dense layer to a dense layer with a single output unit and a sigmoid activation function (for classification) or no activation function (for regression). I tuned the learning rate for training to values of 10^-4^, 10^-3^, or 10^-2^.

The input of the architectures I described thus comprised DNA sequences without any feature engineering. In the architectures below, I used feature engineering that incorporates a simple and generic property of such sequences, namely that individual amino acids are encoded by nucleotide triplets. (I deliberately did not replace codons with the amino acids they encode, because doing so neglects that synonymous codons often have different fitness (Papkou, et al., 2023).) I refrained from using more complex feature engineering, because the resulting features may be specific to my study protein, and because it does not necessarily lead to better performance(Alley, et al., 2019; Asgari and Mofrad, 2015; ElAbd, et al., 2020; Elnaggar, et al., 2021; Iuchi, et al., 2021; Raimondi, et al., 2019; Rao, et al., 2021; Rives, et al., 2021).

##### RNN with codon-based embedding

This architecture is identical to the RNN architecture I described above, except that it used a simple form of feature engineering in which the codons are the elementary input tokens. To this end, I first subdivided the nine nucleotide genotypes into 3 codons (nucleotide triplets), and mapped each codon to an integer between 1 and 64 (in alphabetical order), thus transforming the genotype data into an integer vector of dimension three. I positionally embedded these vectors, assuming an input space of dimension 64 (the number of codons) and an output space whose dimensions I tuned (4, 8, 16, 32). The remaining elements of this architecture, as well as other tuned parameters are identical to that of the RNN architecture above.

##### Transformer with codon-based embedding

As in the RNN above, I first subdivided the nine nucleotide genotypes into 3 codons (nucleotide triplets), and mapped each codon to an integer between 1 and 64 (in alphabetical order), thus transforming each genotype data into an integer vector of dimension three. I positionally embedded these vectors, assuming an input space of dimension 64 (the number of codons) and an output space whose dimensions I tuned (8, 16, 32, 48)

The remainder of this transformer is identical to the above transformer that uses position embedded nucleotide (not codon) data. Specifically, I passed the position-embedded codon data to a multi-head attention layer. I applied residual connection (He, et al., 2016) by adding the output of the embedding layer to the output of the multi-head attention layer, and layer-normalized the resulting data (Xu, et al., 2019). This normalized data then became the input to two consecutive dense layers. I applied residual connection once again by adding the input of the first dense layer to the output of the second dense layer, and then layer-normalized the resulting output.

Collectively, the multi-head attention layer together with the two dense layers, the residual connection, and the layer-normalization constitute one module of the transformer. I then stacked 1, 2, 4, or 6 such modules during tuning. In addition, for each module I tuned the number of attention heads (2, 4, 6 or 8), the key dimension, i.e., the size of each attention head (4, 8, 16, 32, 48), as well as the number of units in each dense layer (4, 8, 16).

Moreover, I tuned whether or not to apply dropout at a fixed rate of 0.1 to the output of the last stack. I flattened the resulting output and fed it into a dense layer with a single output unit and a sigmoid activation function (for classification) or no activation (for regression). I tuned the learning rate, exploring values of 10^-4^, 10^-3^, and 10^-2^.

### Genotype sampling

I explored how different sample sizes and sampling methods of training data affect fitness prediction on test data for three diverse neural network architectures that perform well on the regression task, i.e., the multilayer perceptron, the RNN, and the transformer (the latter two with codon-based positional embedding), focusing mostly on the best-performing RNN (see Supplementary Results 1 for network details). I restricted this analysis to the 17,774 viable genotypes, which encode 1,630 unique amino acid sequences. After choosing suitable training, validation, and test sets as described below. I trained each networks with the rmsprop algorithm for a maximum of 100 epochs with batch sizes of 128 samples. I stopped the training early when the training loss (mean squared error [mse] of fitness prediction) did not decrease for five consecutive epochs. I trained each network in three independent replicates to estimate how much fitness predictions vary across such replicates.

To sample data for each replicate. I first randomly shuffled all genotypes and set aside 50 percent (8887) of them as a test set, and the remainder for validation and training. I then sampled a fixed number of the remaining genotypes for training and validation. I varied this number between *S*=200 (1.1 percent of all data) and *S*=8,000 (45 percent) to explore how prediction quality depends on *S*. (I chose independent test and training/validation data sets for each value of *S* and for each replicate.) Because many of the resulting training/validation data sets were small, I did not use hold-out validation, but applied 4-fold cross-validation, setting aside 75 percent of the sample for training and 25 percent for validation, and repeating this procedure four times with non-overlapping validation data sets for each replicate. I computed the training and validation loss (mse) after each epoch as an average across the four training runs.

Because many of the 1,630 viable amino acid sequences are encoded by multiple nucleotide sequences, a random sample of nucleotide sequences that comprises some fraction *S*/17774 of viable sequences may encode more than the corresponding fraction of 1,630×(*S*/17774) amino acid sequence. For example, a random sample comprising 50 percent of all viable nucleotide sequences encodes on average 1321.4±2.7 (std. err, *n*=10) amino acid sequences, much more than the 0.5×1,630=815 amino acid sequences one might expect. A simple example explains why. Consider a hypothetical data set of 4 nucleotide sequences encoding 2 amino acid sequences, such that 2 nucleotide sequences encode each amino acid sequence.

Now sample 2 of the nucleotide sequences without replacement, and consider a random variable that quantifies that number of amino acid sequences encoded by the nucleotide sequences in this sample. As opposed to the naïve expectation that a sample comprising half of the nucleotide sequences would also encode half of the amino acid sequences, the expectation of this random variable can be computed as (1)×(1/3)+(2)×(2/3)=5/3, because the probability of sampling a second nucleotide sequence encoding the same amino acid sequence as the first sampled sequence equals 1/3. This expected value of 5/3 encoded amino acid sequences is much greater than one, and closer to the maximally possible number of 2 encoded amino acid sequences.

Another consequence of the multiplicity of synonymous nucleotide sequences is that small random samples will be enriched for amino acid sequences that are encoded only by a single nucleotide sequence in the sample. For example, while 34.2% of the 1,630 viable amino acid sequences are encoded by only a single nucleotide sequence, in a small sample of 200 viable nucleotide sequences more than 85 percent of the encoded amino acid sequences are encoded by only a single nucleotide sequence in the sample.

I next describe the methods I used to sample *S* sequences in the training and validation data. In all these methods I still set aside a random (uniform) sample of fifty percent of all viable sequences as a test set, because in most experimental applications, the sequences whose fitness needs to be predicted may be arbitrary and not sampled identically to the training data.

#### Random

I chose all *S* sequences such that each sequence had a uniform probability to be chosen.

#### Fewest synonymous sequences (‘one syn.’)

Here I sampled all sequences at random, but such that each amino acid sequence in the sample is represented only once whenever possible.

That is, among multiple synonymous nucleotide sequences that may encode the same amino acid sequence, I chose only one at random among all such synonymous sequences. For sample sizes *S* larger than the number of unique amino acid sequences in the available data, I first chose one nucleotide sequence for each amino acid sequence at random. I then chose further synonymous DNA sequences at random, such that each amino acid sequences was represented by the smallest possible number of synonymous nucleotide sequences.

#### At least two synonymous sequences per amino acid sequence (‘two syn.’)

Here I sampled all DNA sequences at random, but such that each encoded amino acid sequence in the sample is represented by at least two synonymous DNA sequences if two such DNA sequences exist in the data. Again, because there are many fewer amino acid sequences than nucleotide sequences, it was not possible to meet this criterion exactly for large sample size *S*. In this case, I allowed for more than two synonymous DNA sequences per amino acid sequences, but such that the number of DNA sequences per amino acid sequence beyond two is as small as possible. In addition, a substantial fraction of amino acid sequences (34.2% of the 1,630 viable amino acid sequences) is encoded by only a single nucleotide sequence. I included such sequences in this sampling procedure by sampling the only nucleotide sequence encoding them, because excluding them would bias my sampling against such sequences.

#### Maximally diverse nucleotide sequence (‘DNA div.’)

Here I sampled DNA sequences such that they are maximally diverse in terms of their pairwise sequence distance. I quantify this distance as the pairwise Hamming distance, i.e., the number of single nucleotide changes that are minimally needed to convert one sequence into the other. To achieve this sampling, I used a heuristic procedure, for which I first numerically encoded each DNA sequence as a flattened one-hot encoded vector of dimension 9×4=36. I then sampled a sequence at random. Subsequently, I chose among all remaining sequences one that had the maximal distance from the sampled sequence and added it to the sample. In the sequence encoding I use, each sequence can be viewed as a point or location in a 36-dimensional space. I computed the ‘average location’ of the two already sampled sequences by averaging their one-hot encoded vector representation. (More generally, for any number of sequences, this average can also be viewed as the center of mass of the sequences in 36-dimensional sequence space.) For all subsequent sequences to be sampled I proceeded analogously. That is, I sampled a sequence that had the maximum distance from the center of mass of the already sampled sequences, and added it to the sample until I had sampled all *S* sequences.

#### Maximally (physicochemically) diverse amino acid sequences (‘protein div.’)

Amino acids can be distinguished by physicochemical features that can help predict the effect of amino acid changes on a protein’s phenotype. Here I used a categorization of amino acids that distils more than 500 amino acid features into 19 properties that are maximally independent of each other (Georgiev, 2009). I chose this representation because it outperforms others in similar machine learning tasks (Wittmann, et al., 2021). I obtained data for this representation from computer code in (Ofer and Linial, 2015). I represented each amino acid as a 19-dimensional vector that quantifies these properties, which resulted in a 3×19=57-dimensional vector representation for each amino acid sequence in my data set.

I then sampled *S* DNA sequences that are maximally diverse with respect to the physicochemical properties of the encoded amino acid sequences. To create this maximally diverse data set, I used the same procedure as in the previous section on sampling maximally diverse DNA sequences, except that now I sampled DNA sequences that are maximally distant from each other based on the Euclidean distance of their encoded amino acid sequences in the 57-dimensional space.

#### Maximal codon usage (‘CUB’)

Here I preferentially sampled DNA sequences in which the amino acids are encoded by preferred codons, i.e., codons that are found in highly expressed proteins and that facilitate translation, because their cognate tRNAs are highly abundant (Hershberg and Petrov, 2008; Komar, 2016). To this end, I obtained data on codon usage in *E.coli* strain *K12* from the Kazusa codon usage data base (Nakamura, et al., 2000) (https://www.kazusa.or.jp/codon/), which reports the average number of codons encoding a specific amino acid among 1,000 codons. Using this data, I then ranked, for each amino acid, the codons encoding the amino acid from most preferred to least preferred. (I analogously ranked stop codons by the frequency of their occurrence.) I then computed for each nucleotide sequence the sum of the preference scores for the encoded amino acids (and stop codons). Subsequently, I catalogued for each amino acid sequence the set of encoding nucleotide sequences, and ranked these nucleotide sequences by their codon preference score. If the sample size *S* was smaller than the total number of viable amino acid sequences, I then chose a random subset of these amino acid sequences such that each sequence was encoded by the most preferred nucleotide sequence for that score. If the sample size was larger, I first chose an amino acid sequence at random, and then the encoding nucleotide sequence with the highest preference score. I repeated this procedure (without replacement of nucleotide sequences) until I had sampled *S* nucleotide sequences.

#### Conventional ‘codon compression’ sampling strategies

I randomly sampled *S* nucleotide sequences (with uniform probability) from the subset of sequences where (i) all third codon positions had either a G or T in the third position (NNK sampling, K=G/T), (ii) all third codon positions had either a C or G in the third position (NNS sampling, *S*=C/G), (iii) all third codon positions had a T in the third position (NNT sampling), (iv) all third codon positions had a G in the third position (NNG sampling), (v) all second codon positions had A, G, or T (A/G/T=D), and all third codon positions had a T (NDT sampling).

## Supplementary Results 1

### Multiple deep learning architectures can identify viable genotypes with high sensitivity and specificity

Mutagenesis of typical proteins creates mostly non-functional variants(Li, et al., 2019; Wittmann, et al., 2021; Wu, et al., 2019). DHFR is no exception, because only a small minority (6.8 percent) of genotypes in its trimethoprim resistance landscape is viable(Papkou, et al., 2023). Thus, the most basic machine learning task is to distinguish viable from inviable genotypes with a binary classifier. To quantify the performance of any such classifier I use both sensitivity – the probability of correctly predicting the viability of an actually viable genotype – and specificity – the probability of correctly predicting the inviability of an actually inviable genotype (Methods).

I first computed these performance measures for two simple benchmarks. The first is a random predictor, for which I randomly shuffled the measured fitness value among all genotypes. This predictor has a low sensitivity of 0.07 to predict viable genotypes. In contrast, it has a high specificity of 0.93 to predict inviable genotypes. This is not surprising, given that most genotypes are inviable (Table S1). The second benchmark is a predictor based on linear regression (Methods), for which I used a flattened one-hot encoded representation of the genotype as a 9×4=36-dimensional variable. This linear predictor is much more sensitive (0.82) than the random predictor, but no more specific (Table S1).

I next studied the classification performance of four fundamental deep learning neural network architectures. The first architecture is the multilayer perceptron (Gurney, 1997; LeCun, et al., 2015; Rosenblatt, 1958), in which I tuned the number of layers, the number of neurons per layer, weight regularization, layer dropout, and the learning rate (Methods). The best-performing such network improved its sensitivity by 10.5 percent and its specificity by 7.4 percent relative to linear regression (Table S1). The second architecture is a bidirectional recurrent neural network (Hochreiter and Schmidhuber, 1997), in which I tuned the number of bidirectional layers, the number of neurons in each layer, weight regularization, recurrent dropout, and the learning rate. This architecture performed better than the multilayer perceptron, yielding an improvement of 14.0 percent in sensitivity relative to linear regression. The improvement in specificity (7.2 percent) was similar to that of the perceptron. More generally, because this and all architectures below showed high specificity (>0.996, Table S1), I will focus on sensitivity as a measure of performance improvement. The third architecture was a one-dimensional convolutional network (LeCun, et al., 2015), in which I tuned the number of convolutional layers, the number of dense layers that followed them, the number of neurons in these layers, their weight regularization, and the learning rate. This classifier performed slightly worse than the recurrent network, with an improvement of 13.4 percent relative to linear regression.

I deliberately chose such end-to-end learning of word embedding, because it performs at a par with highly complex pre-trained embeddings, may require lower embedding dimensions, and does not depend on other bioinformatic resources (Alley, et al., 2019; Asgari and Mofrad, 2015; ElAbd, et al., 2020; Elnaggar, et al., 2021; Iuchi, et al., 2021; Raimondi, et al., 2019; Rao, et al., 2021; Rives, et al., 2021). In this transformer architecture, I tuned the number of embedding dimensions, the number of attention heads per transformer module, the size of each attention head, the number of neurons in each dense layer of a module, the number of stacked transformer modules, the dropout rate, and the learning rate (Methods).

The best transformer performed worse than the recurrent network (sensitivity improvement 10.7 percent relative to linear regression), and not much better than the multilayer perceptron (10.5 percent).

Feature engineering, i.e., choosing an appropriate representation of input data, can be crucial to improve network performance (Chollet, 2021). For two further neutral networks, I chose a simple and general form of feature engineering that would apply to all protein-coding genes and is not specific to DHFR or a specific protein class. Specifically, I subdivided the 9 nucleotide input sequence into 3 integer-encoded codons and positionally embedded these codons into a space whose dimensionality I varied during hypertuning (Methods). These codons became the input to a bidirectional RNN whose hyperparameters I also tuned (Methods). The best performing RNN showed a sensitivity improvement of 13.2% relative to linear regression (Table S1). I then also used the same codon-based learned embedding as an input to a transformer that I tuned as just described for the simpler transformer architecture (Methods). Here the best network was 13.9% more sensitive than linear regression. Thus, in neither network does feature engineering improve the sensitivity of the best architecture without it (RNN, sensitivity 0.935), although the differences are small – less than 1% in sensitivity and identical specificity (Table S1).

In sum, these data show that even architectures as simple as a multilayer perceptron can identify viable genotypes with both high sensitivity and specificity, a performance that can be further improved by more complex architectures, most notably recurrent neural networks.

Not surprisingly given the differences of the tasks, the values of the hyperparameters – including the total number of parameters – associated with the best-performing networks differ between binary classification and fitness prediction (Supplementary Results 2, Table S1 and Table 1).

## Supplementary Results 2

The following sections describe the best-performing networks after hypertuning. See methods for details on tuned hyperparameters.

### Binary classification

#### 1.1 Multilayer perceptron

Three dense layers of 32 output units each, followed by one dense layer with a single unit and sigmoid activation, residual connection between output of layer one and three, layer normalization after layer 3, no dropout, learning rate 10^-3^, 3,393 parameters.

#### 1.2. Recurrent neural network

4 bidirectional (LSTM) recurrent neural network layers with 48 units each, followed by one dense layer with a single unit and sigmoid activation function, residual connection between the output of recurrent layers 1 and 2, 2 and 3; batch normalization of output of layer 2 and 3 (after residual connection), recurrent layer dropout of 0.1, learning rate 10^-3^, 188,257 parameters.

#### 1.3. Transformer

8 embedding dimensions, 6 stacked transformer modules, each with 8 attention heads, 4 key dimensions (the size of each attention head), and 4 units in the dense layers, dropout of 0.1 after the last module, followed by one dense layer with a single unit and sigmoid activation, learning rate 10^-3^,8829 parameters.

#### 1.4. Convolutional network

3 convolutional layers (with 24, 32, and 48 filters) followed by two dense layers with 32 units each, no regularization, learning rate of 10^-3^, 9,769 parameters.

#### 1.5. RNN with codon-based positional embedding

8 embedding dimensions, 4 bidirectional (LSTM) recurrent neural network layers with 32 units each, followed by one dense layer with a single unit, residual connection between the output of recurrent layers 1 and 2, 2 and 3; batch normalization of layer 2 and 3 output (after residual connection), no dropout, no regularization, learning rate 10^-3^, 86,105 parameters.

#### 1.6. Transformer with codon-based positional embedding

32 embedding dimensions, 5 stacked transformer modules, each with 2 attention heads, 16 key dimensions (the size of each attention head), and 8 units in the dense layers of each transformer module, followed by one dense layer with a single unit, no dropout for any layer, no regularization, learning rate 10^-3^, 26,761 parameters.

### Regression

#### 2.1 Multilayer perceptron

7 dense layers of 32 output units each, followed by one dense output layer with a single unit, layer normalization after layer 3, 5, and 7, residual connection between output of layers 1 and 3, 3 and 5 (after layer normalization), as well as 5 and 7 (after layer normalization), no dropout, no regularization, learning rate 10^-2^, 7,745 parameters (see also Figure S12).

#### 2.2. Recurrent neural network

5 bidirectional (LSTM) recurrent neural network layers with 16 units each, followed by one dense layer with a single unit, residual connection between the output of recurrent layers 1 and 2, 2 and 3, 3 and 4; batch normalization of layer 2, 3, and 4 output (after residual connection), no recurrent layer dropout, learning rate 10^-2^, 28,193 parameters.

#### 2.3. Transformer

8 embedding dimensions, 6 stacked transformer modules, each with 8 attention heads, 2 key dimensions (the size of each attention head), and 4 units in the dense layers of each transformer module, no dropout, last transformer followed by one dense layer with a single unit, learning rate 10^-3^, 4,909 parameters.

#### 2.4. Convolutional network

5 convolutional layers (with 24, 32, 48, 64, and 96 filters) followed by two dense layers with 16 units each, learning rate of 10^-2^, 16,233 parameters.

#### 2.5. RNN with codon-based positional embedding

32 embedding dimensions, 5 bidirectional (LSTM) recurrent neural network layers with 16 units each, followed by one dense layer with a single unit, residual connection between the output of recurrent layers 1 and 2, 2 and 3, 3 and 4, batch normalization of layer 2, 3, and 4 output (after residual connection), recurrent layer dropout of 0.1, learning rate 10^-2^, 33,921 parameters. (see also Figure S13).

#### 2.6. Transformer with codon-based positional embedding

32 embedding dimensions, 6 stacked transformer modules, each with 8 attention heads, 16 key dimensions (the size of each attention head), and 16 units in the dense layers of each transformer module, dropout of 0.1 after the last module, followed by one dense layer with a single unit, learning rate 10^-3^,128,241 parameters (see also Figure S14).

## Supplementary Results 3

Here I ask how simple and widely used conventional codon compression sampling methods influence fitness predictions at different training sample sizes *S* compared to random sampling(Pines, et al., 2015). Combinatorial libraries of genotypes are usually built with degenerate synthetic oligonucleotides that admit multiple nucleotides at individual nucleotide positions. Because synthesizing such oligonucleotides can be costly for complex patterns of degeneracy, simpler codon compression strategies than I analyzed in the main text are often used. Most prominent among them is ‘NNT’ sampling, where the first and second position of each codon admit any nucleotide but the third position is restricted to a T; ‘NNG’ sampling, where the third position is restricted to a G; ‘NNK’ sampling (third position: G or T), ‘NNS’ sampling (third position: C or G); or ‘NDT sampling (second position: A, G or T; third position: T). These strategies dramatically reduce the complexity of a library. For example, in a combinatorially complete library of three codons that harbor all 4^9^ possible nucleotide sequences, NNT sampling reduces the number of sequences in the library by a factor 4^3^=64. Unfortunately, this dramatic reduction in library size also means that for most of these strategies, the present data set contains too few viable sequences to evaluate prediction performance. Specifically, among all 17,774 viable sequences, only 327, 259, 2367, 2442, and 186 fulfill the criteria imposed by NNT, NNG, NNK, NNS, and NDT sampling.

Thus, for three of the five sampling procedures too few sequences exist to evaluate prediction performance. The exceptions are NNK and NNS sampling, where it is possible to evaluate the performance of a training/validation sample comprising of the order of *S*=1,000 sequences. For the RNN network at *S*=1,000, NNK sampling yields a mape of 20.9±1.18 compared to random sampling at 18.7±5.8, and an *R^2^*=0.17 compared to *R^2^*=0.65. (perceptron: mape=12.4±0.35 vs. 13.1±1.26, *R^2^*=0.58 vs. 0.59 for random sampling; transformer: mape=17.7±0.57 vs. 11.7±0.11, *R^2^*=0.21 vs. 0.64 for random sampling, see also Figures S5, S6, and S7) Also for the RNN network at *S*=1,000, NNS sampling yields a mape of 23.7±1.27 compared to random sampling at 18.7±5.8, and an *R^2^*=0.16 compared to *R^2^*=0.65. (perceptron: mape=12.5±0.63 vs. 13.1±1.26, *R^2^*=0.57 vs. 0.59 for random sampling; transformer: mape=20.2±1.47 vs. 11.7±0.11, *R^2^*=0.17 vs. 0.64 for random sampling, Figures S5, S6, and S7)

In sum, for the RNN architecture, the mape is very similar between NNK sampling, NNS sampling, and random sampling. In contrast, NNK and NNS sampling both reduce the percent of explained variation in fitness (*R^2^*) dramatically. For the multilayer perceptron, both procedures perform similarly to random sampling with respect to both mape and *R^2^*, whereas for the transformer, they perform worse with respect to both mape and *R^2^*.

## Supplementary Tables

**Table S1:** Performance of deep learning network architectures on binary classification.

| | sensitivity | specificity | CE<sup>a</sup> | parameters ($\times 10^3$)<sup>b</sup> |
| --- | --- | --- | --- | --- |
| Random | 0.07 | 0.93 | n/a | n/a |
| Linear regression | 0.82 | 0.93 | n/a | n/a |
| Perceptron | 0.906 (10.5%)<sup>c</sup> | 0.999 (7.4%) | 0.030 | 3.4 |
| RNN | 0.935 (14.0%) | 0.997 (7.2%) | 0.032 | 188.9 |
| Convolutional | 0.930 (13.4%) | 0.998 (7.3%) | 0.027 | 9.8 |
| Transformer | 0.908 (10.7%) | 0.997 (7.2%) | 0.035 | 8.8 |
| RNN (codons)<sup>d</sup> | 0.928 (13.2%) | 0.997 (7.2%) | 0.029 | 86.1 |
| Transformer (codons)<sup>d</sup> | 0.934 (13.9%) | 0.997 (7.2%) | 0.030 | 26.8 |

<sup>a</sup> CE: binary cross-entropy, loss function used for neural network training. <sup>b</sup> number of parameters in the best performing architecture. <sup>c</sup> Numbers in parentheses indicate percent improvement relative to linear regression. <sup>d</sup> Architecture with codon-based positional embedding.

## Supplementary Tables

**Figure S1.**
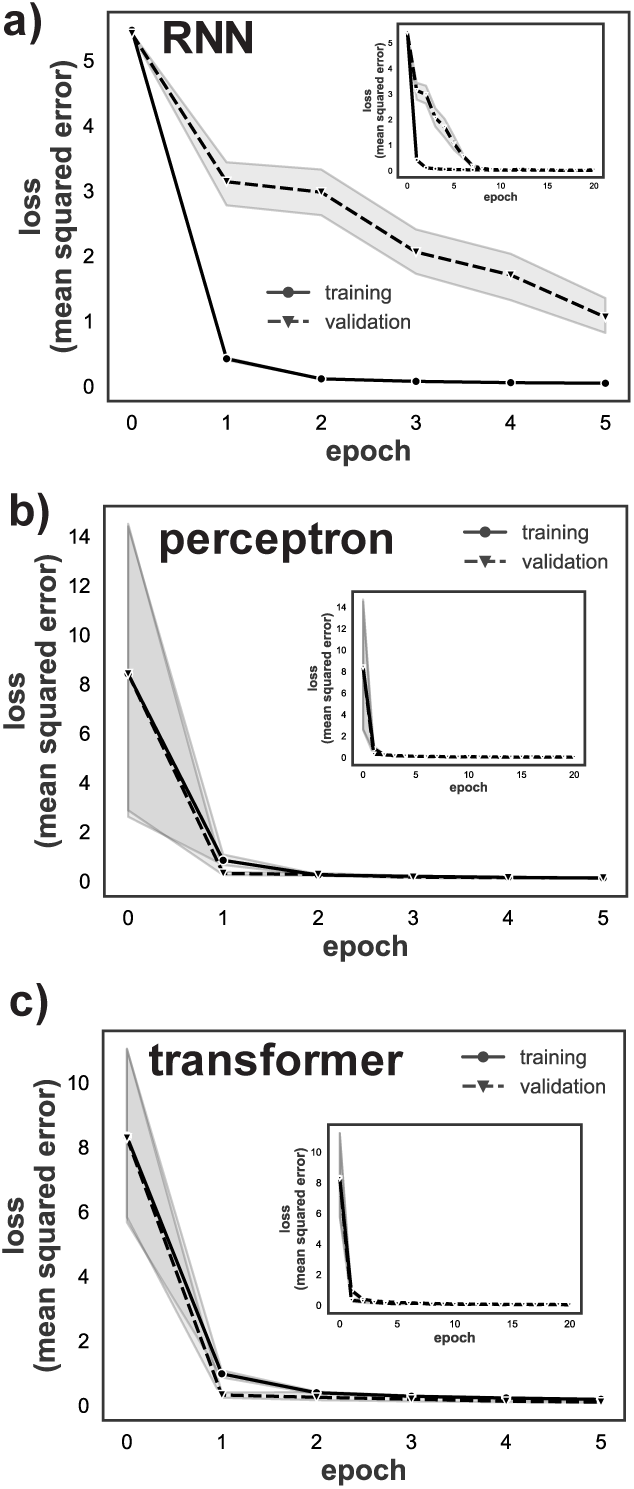
Training and validation loss of the three best-performing neural network architectures during training. Training and validation loss (mean squared error, vertical axis) of the best-performing **a)** recurrent neural network (RNN) using positional embedding of codons, **b)** multilayer perceptron, and **c)** transformer using positional embedding of codons (See Supplementary Results 1 for architecture details) as a function of training time (epochs, horizontal axis). Insets show the same data but for 20 epochs. Training loss declines rapidly and approaches values close to its minimum within 10 epochs for all architectures. A lack of overfitting for the training data even at later times indicates that the hypertuned architectures could still be substantially improved. Data in each panel is based on 5 identical replicate trainings starting from the same initial architecture. Symbols connected by lines and shaded areas indicates mean loss and 95% confidence intervals, respectively, based on these five replicates. Absent shading indicates that confidence intervals are too small to be visible. I chose the training and validation data for each panel identical to that used during hypertuning (methods). That is, I split the 17774 viable genotypes into a training and a validation data set that comprised 25 percent of these genotypes each, and for which I sampled genotypes randomly and without replacement. Like elsewhere, I used the rmsprop algorithm for gradient descent, with minibatch sizes of 128 samples.

**Figure S2:**
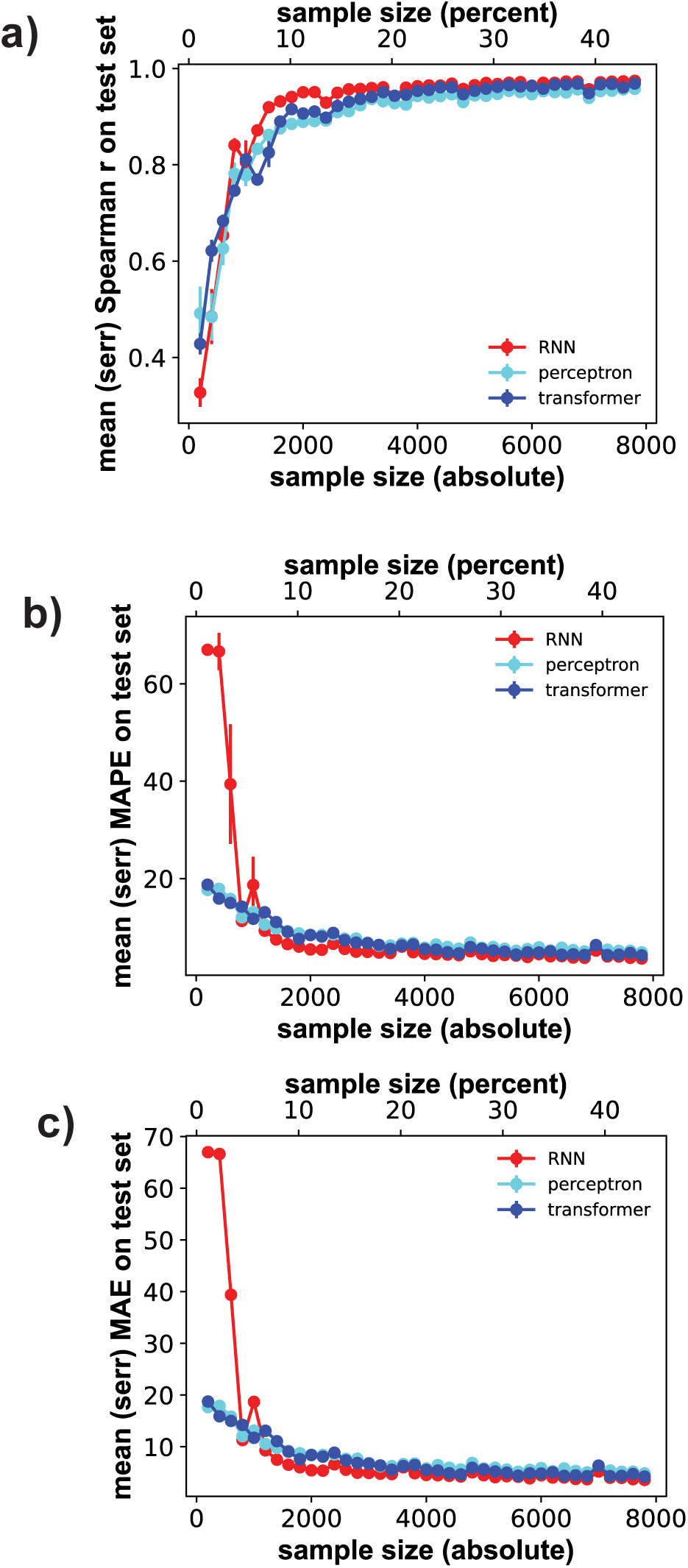
Predictive performance increases rapidly with sample size for all neural network architectures. The horizontal axis of all panels shows the size *S* of the genotype sample used for training and validation through 4-fold cross-validation, both in absolute numbers of genotypes (bottom) and as a percentage of all viable genotypes (top). The vertical axes show the performance of the three major network architectures (color legend) on a test-set comprising 50% (8887) of viable genotypes, as quantified by **a)** the Spearman correlation coefficient between measured and predicted fitness, **b)** the minimum absolute percentage error (mape), and **c)** the minimum absolute error (mape) of predicted fitness. I chose genotypes for training, validation, and testing genotypes randomly and with uniform probability among all viable genotypes.

**Figure S3:**
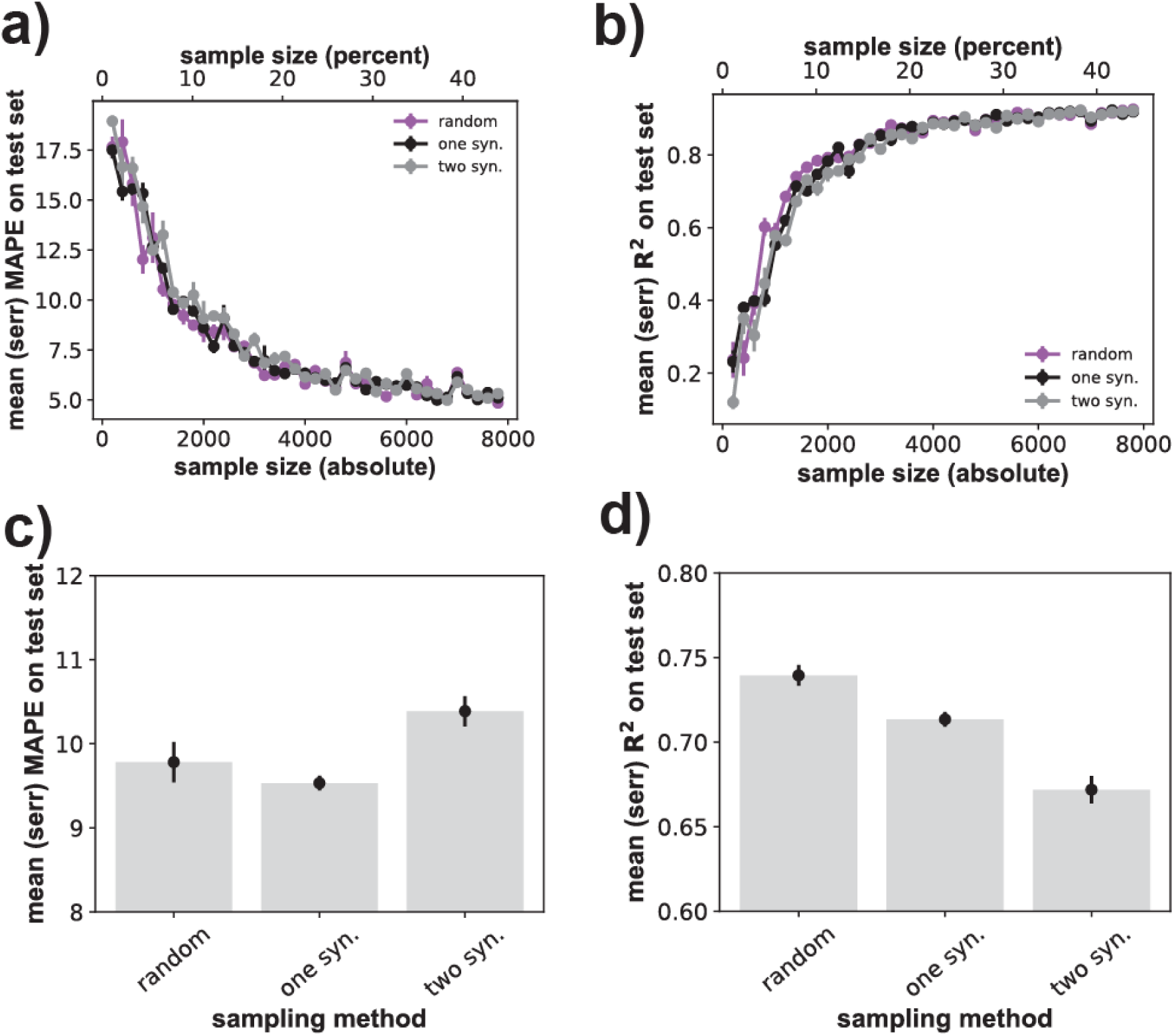
Sampling one or two synonymous sequences does not strongly affect multilayer perceptron prediction quality. **a)** The horizontal axis shows the size *S* of the genotype sample used for training and validation through 4-fold cross-validation, both in absolute numbers of genotypes (bottom) and as a percentage of all viable genotypes (top). The vertical axis shows the prediction quality of the best-performing multilayer perceptron architecture, as quantified by the mape of fitness prediction as a function of sample size *S*. The *S* genotypes are either sampled randomly and uniformly (‘random’), or such that only one synonymous (‘one syn.’) or two synonymous (‘two syn.’) nucleotide sequences are sampled per amino acid sequence (Methods). Whiskers indicate one standard error of the mean based on three replicate trainings for each network and sample size. **b)** like a), but prediction quality is quantified through the coefficient of determination *R^2^*. **c)** Dot-whisker plot indicating the means (height of bars) and standard errors (whiskers) of the mape at a fixed sample size of *S*=1,400 genotypes for the three sampling methods shown on the horizontal axis. **d)** like c), but for *R^2^*instead of the mape.

**Figure S4:**
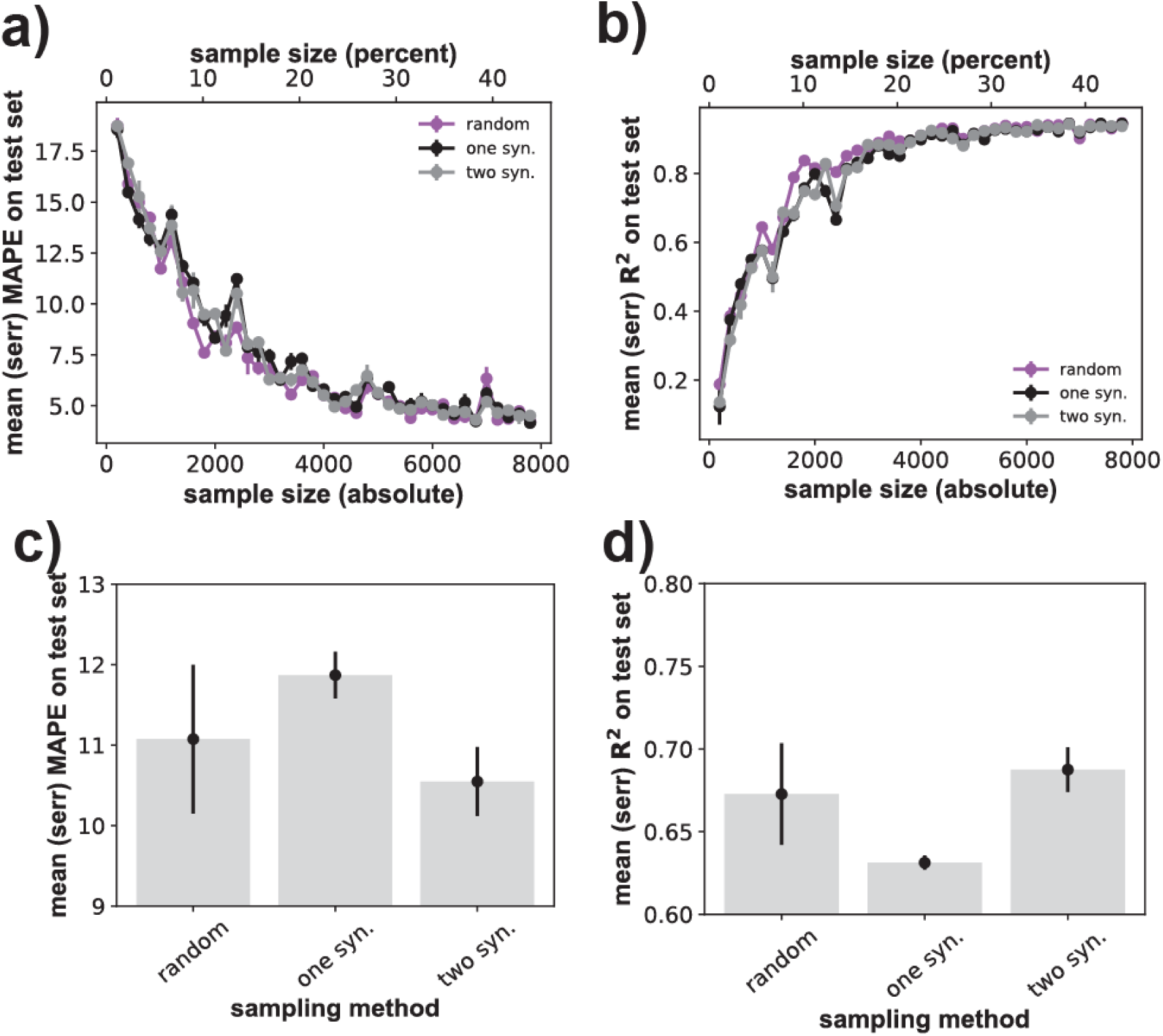
Sampling one or two synonymous sequences does not strongly affect transformer prediction quality. **a)** The horizontal axis shows the size *S* of the genotype sample used for training and validation through 4-fold cross-validation, both in absolute numbers of genotypes (bottom) and as a percentage of all viable genotypes (top). The vertical axis shows the prediction quality of the best-performing transformer architecture, as quantified by the mape of fitness prediction as a function of sample size *S*. The *S* genotypes are either sampled randomly and uniformly (‘random’), or such that only one synonymous (‘one syn.’) or two synonymous (‘two syn.’) nucleotide sequences are sampled per amino acid sequence (Methods). Whiskers indicate one standard error of the mean based on three replicate trainings for each network and sample size. **b)** like a), but prediction quality is quantified through the coefficient of determination *R^2^*. **c)** Dot-whisker plot indicating the means (height of bars) and standard errors (whiskers) of the mape at a fixed sample size of *S*=1,400 genotypes for the three sampling methods shown on the horizontal axis. **d)** like c), but for *R^2^* instead of the mape.

**Figure S5:**
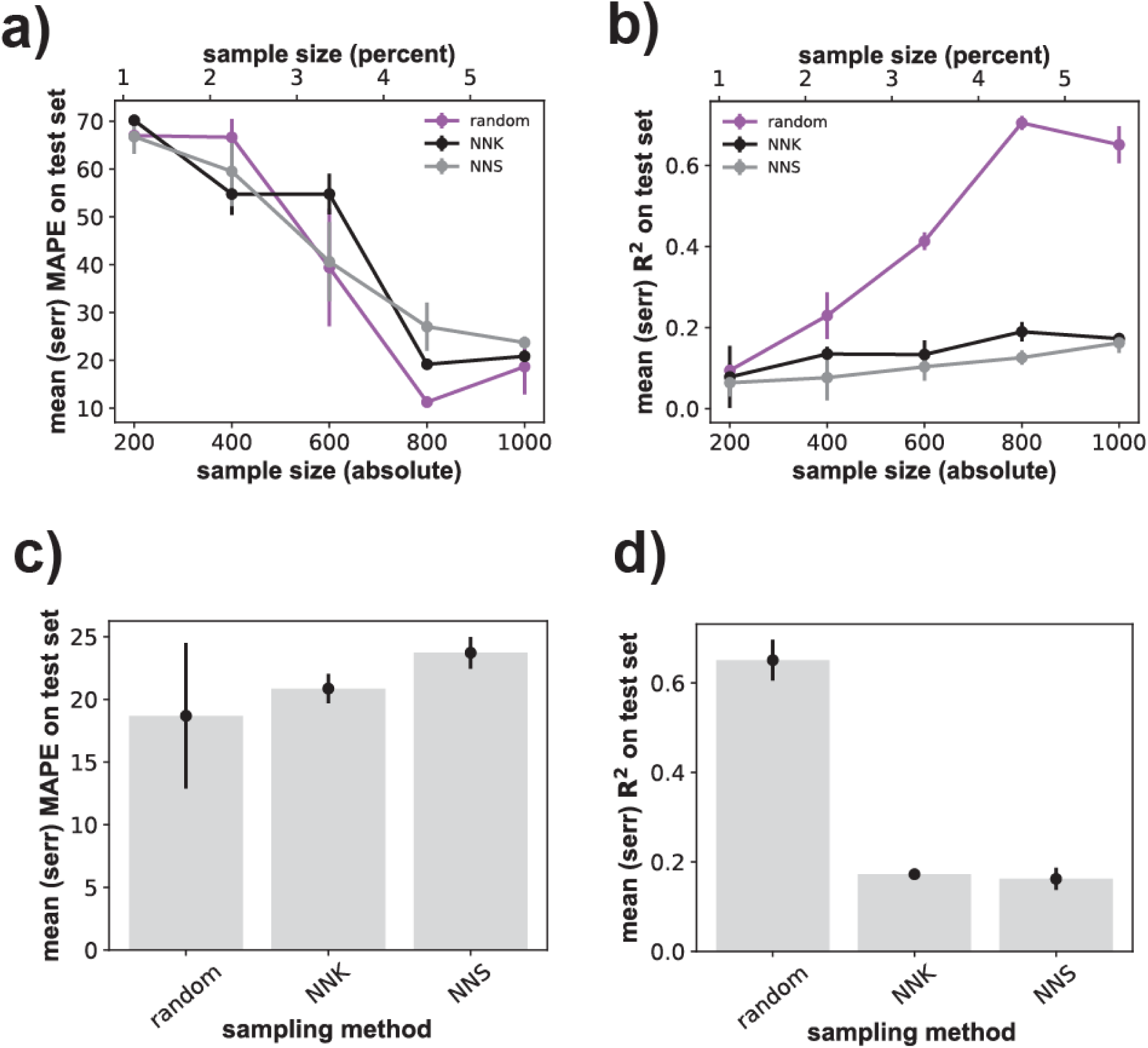
NNS and NNK codon compression genotype sampling substantially reduces the *R^2^*(but not the mape) of the RNN. **a)** The horizontal axis shows the size *S* of the genotype sample used for training and validation through 4-fold cross-validation, both in absolute numbers of genotypes (bottom) and as a percentage of all viable genotypes (top). The vertical axis shows the fitness prediction performance (mape) of the best-performing RNN architecture, as quantified by the mape of fitness predictions as a function of sample size *S*. The *S* genotypes are either sampled randomly and uniformly (‘random’), or such that at each codon’s third position only the nucleotides G/T (‘NNK’) or C/G (‘NNS’) are allowed (Methods). Only sample sizes up to *S*=1,000 genotypes are considered here, because NNK and NNS sampling severely restrict the number of admissible genotypes, rendering much larger training/validation samples infeasible. Whiskers indicate one standard error of the mean based on three replicate trainings for each network and sample size. **b)** like a), but prediction quality is quantified through the coefficient of determination *R^2^*. **c)** Dot-whisker plot indicating the means (height of bars) and standard errors (whiskers) of the mape at a fixed sample size of *S*=1,000 genotypes for the three sampling methods shown on the horizontal axis. **d)** like c), but for *R^2^*instead of the mape.

**Figure S6:**
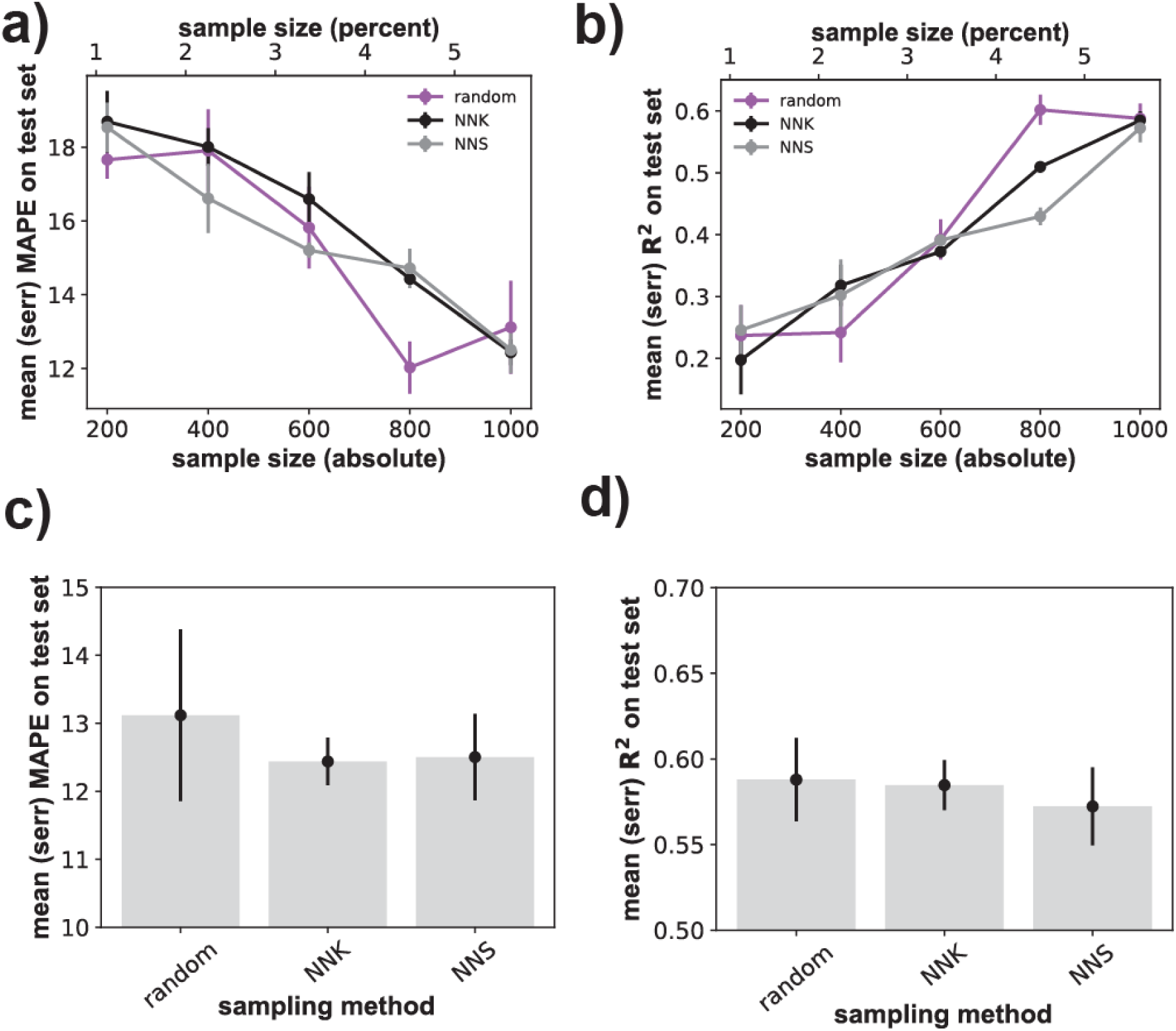
NNS and NNK codon compression genotype sampling do not affect prediction quality of the multilayer perceptron substantially. **a)** The horizontal axis shows the size *S* of the genotype sample used for training and validation through 4-fold cross-validation, both in absolute numbers of genotypes (bottom) and as a percentage of all viable genotypes (top). The vertical axis shows the prediction quality of the best-performing multilayer perceptron architecture, as quantified by the mape of predicted fitness as a function of sample size *S*. The *S* genotypes are either sampled randomly and uniformly (‘random’), or such that at each codon’s third position only the nucleotides G/T (‘NNK’) or C/G (‘NNS’) are allowed (Methods). Only sample sizes up to *S*=1,000 viable genotypes are considered here, because the NNK and NNS sampling severely restrict the number of admissible genotypes, rendering much larger training/validation samples infeasible. Whiskers indicate one standard error of the mean based on three replicate trainings for each network and sample size. **b)** like a), but prediction quality is quantified through the coefficient of determination *R^2^*. **c)** Dot-whisker plot indicating the means (height of bars) and standard errors (whiskers) of the mape at a fixed sample size of *S*=1,000 genotypes for the three sampling methods shown on the horizontal axis. **d)** like c), but for *R^2^* instead of the mape.

**Figure S7:**
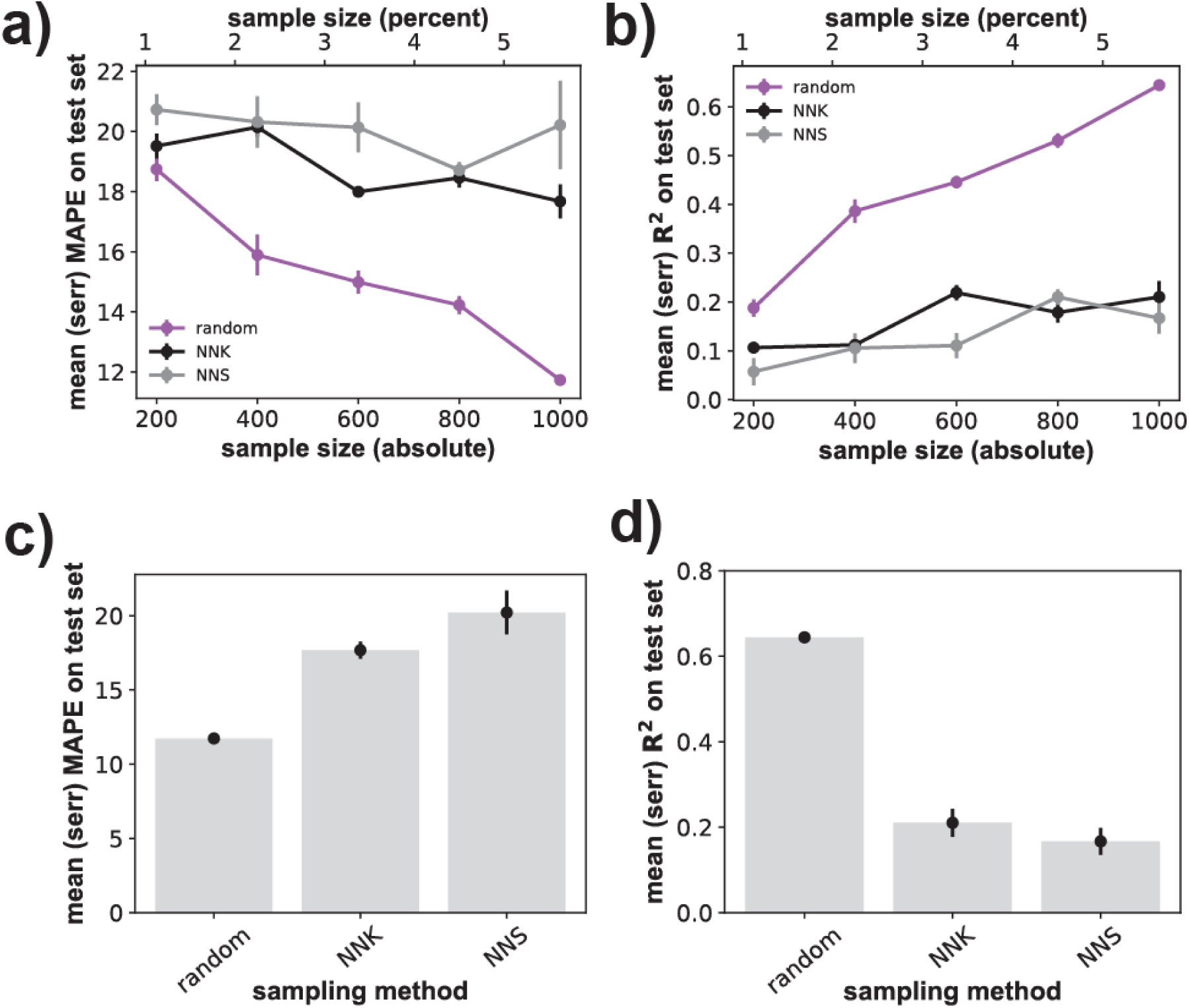
NNS and NNK ‘codon compression’ genotype sampling substantially reduce the prediction quality of the transformer architecture. **a)** The horizontal axis shows the size *S* of the genotype sample used for training and validation through 4-fold cross-validation, both in absolute numbers of genotypes (bottom) and as a percentage of all viable genotypes (top). The vertical axis shows the prediction quality of the best-performing transformer architecture, as quantified by the mape of predicted fitness as a function of sample size *S*. The *S* genotypes are either sampled randomly and uniformly (‘random’), or such that at each codon’s third position only the nucleotides G/T (‘NNK’) or C/G (‘NNS’) are allowed (Methods). Only sample sizes up to *S*=1,000 viable genotypes are considered here, because NNK and NNS sampling severely restrict the number of admissible genotypes, rendering much larger training/validation samples infeasible. Whiskers indicate one standard error of the mean based on three replicate trainings for each network and sample size. **b)** like a), but prediction quality is quantified through the coefficient of determination *R^2^*. **c)** Dot-whisker plot indicating the mean (height of bars) and standard error (whiskers) of the mape at a fixed sample size of *S*=1,000 genotypes for the three sampling methods shown on the horizontal axis. **d)** like c), but for *R^2^* instead of the mape.

**Figure S8:**
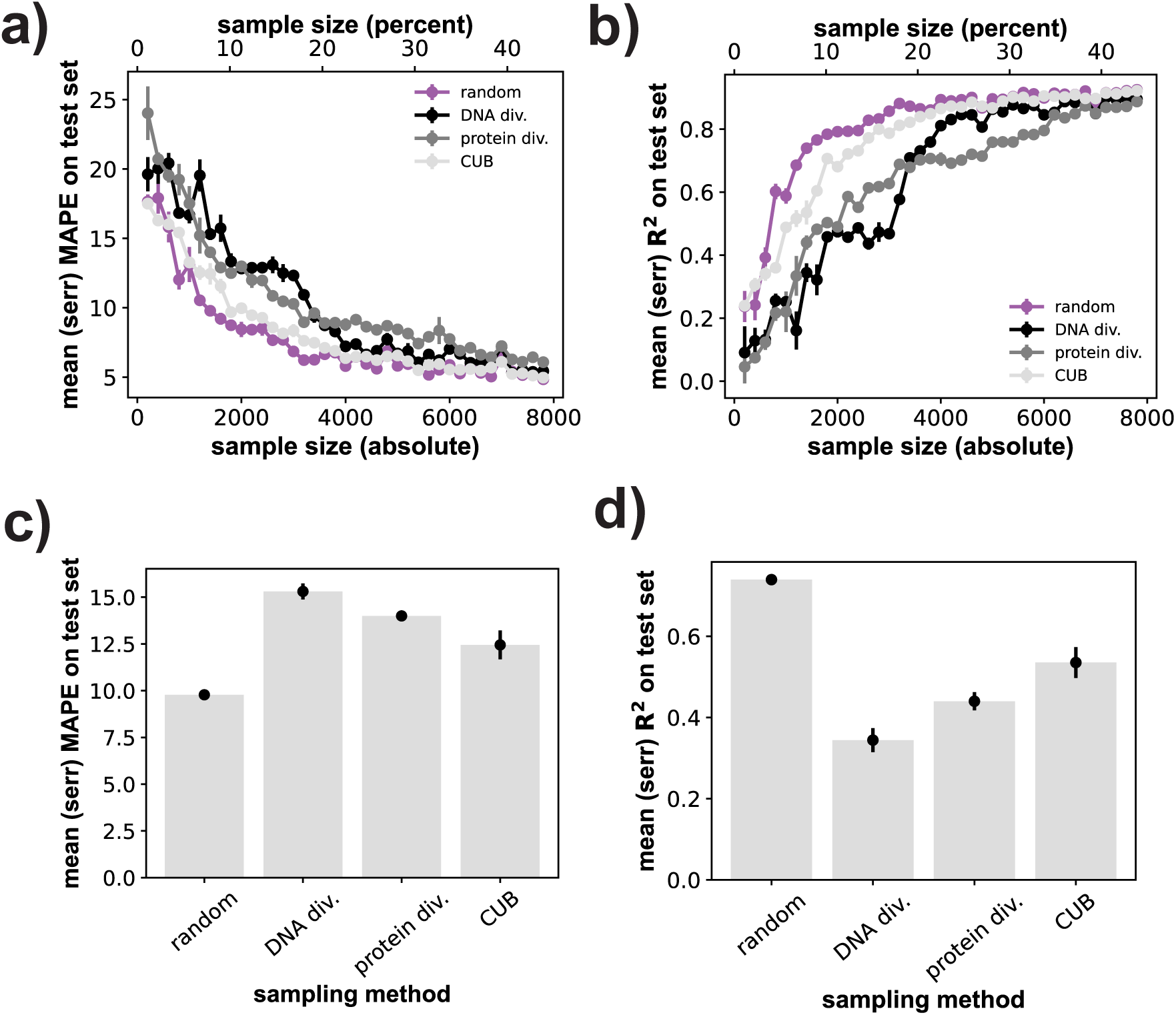
Sampling for sequence diversity or codon usage bias substantially reduces multilayer perceptron prediction quality. **a)** The horizontal axis shows the size *S* of the genotype sample used for training and validation through 4-fold cross-validation, both in absolute numbers of genotypes (bottom) and as a percentage of all viable genotypes (top). The vertical axis shows the prediction quality of the best-performing multilayer perceptron architecture, as quantified by the mape of predicted fitness as a function of sample size *S*. The *S* genotypes are either sampled randomly and uniformly (‘random’), to achieve maximal DNA sequence diversity (‘DNA div.’), maximal amino acid sequence diversity (‘protein dev.’), or maximal codon usage bias (‘CUB’, Methods). Whiskers indicate one standard error of the mean based on three replicate trainings for each network and sample size. **b)** like a), but prediction quality is quantified through the coefficient of determination *R^2^*. **c)** Dot-whisker plot indicating the means (height of bars) and standard errors (whiskers) of the mape at a fixed sample size of *S*=1,400 genotypes for the three sampling methods shown on the horizontal axis. **d)** like c), but for *R^2^* instead of the mape.

**Figure S9:**
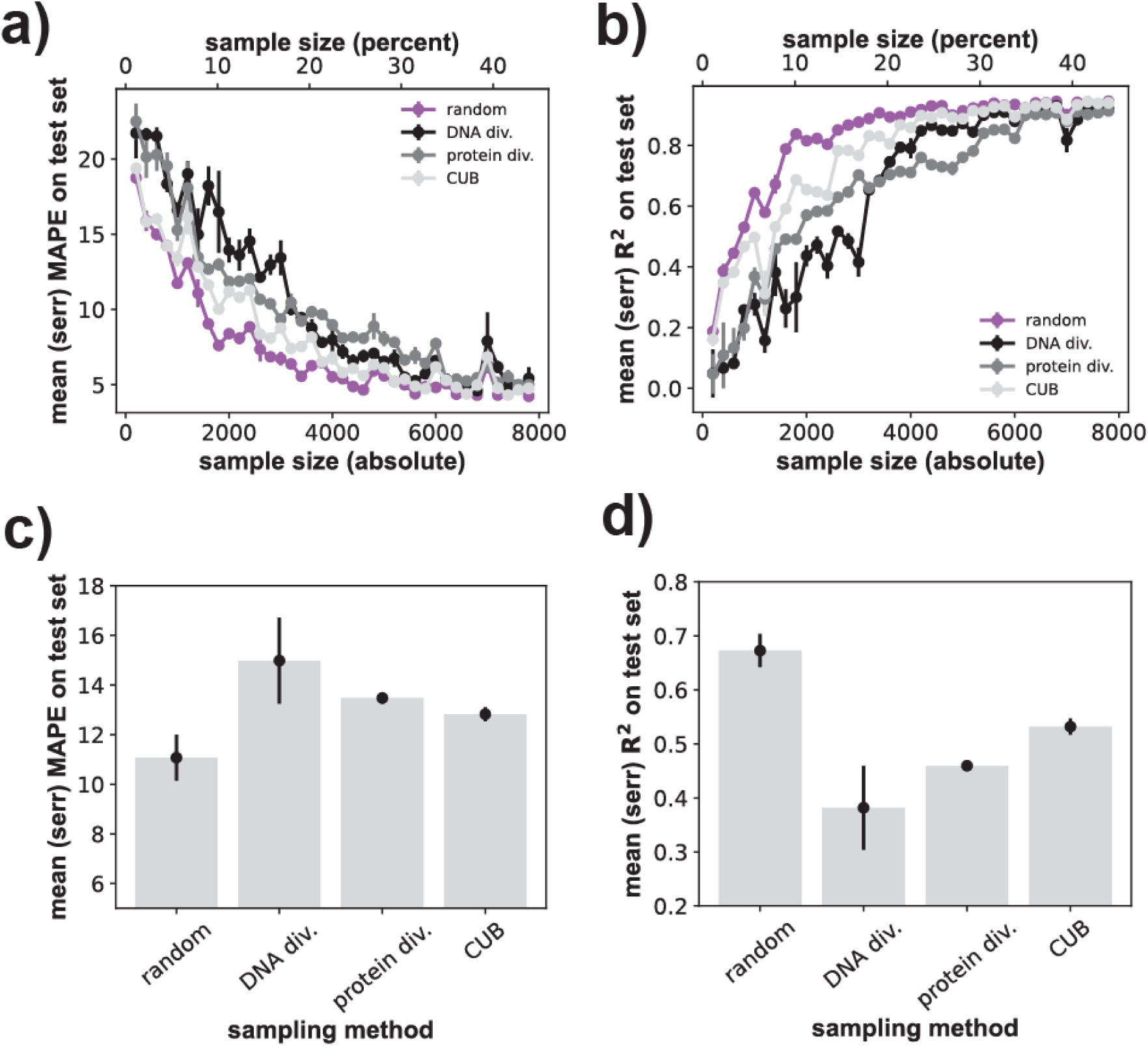
Sampling for sequence diversity or codon usage bias substantially reduces transformer prediction quality. **a)** The horizontal axis shows the size *S* of the genotype sample used for training and validation through 4-fold cross-validation, both in absolute numbers of genotypes (bottom) and as a percentage of all viable genotypes (top). The vertical axis shows the prediction quality of the best-performing transformer architecture, as quantified by the mape of predicted fitness as a function of sample size *S*. The *S* genotypes are either sampled randomly and uniformly (‘random’), to achieve maximal DNA sequence diversity (‘DNA div.’), maximal amino acid sequence diversity (‘protein dev.’), or maximal codon usage bias (‘CUB’) (Methods). Whiskers indicate one standard error of the mean based on three replicate trainings for each network and sample size. **b)** like a), but prediction quality is quantified through the coefficient of determination *R^2^*. **c)** Dot-whisker plot indicating the means (height of bars) and standard errors of the mean (whiskers) of the mape at a fixed sample size of *S*=1,400 genotypes for the three sampling methods shown on the horizontal axis. **d)** like c), but for *R^2^* instead of the mape.

**Figure S10:**
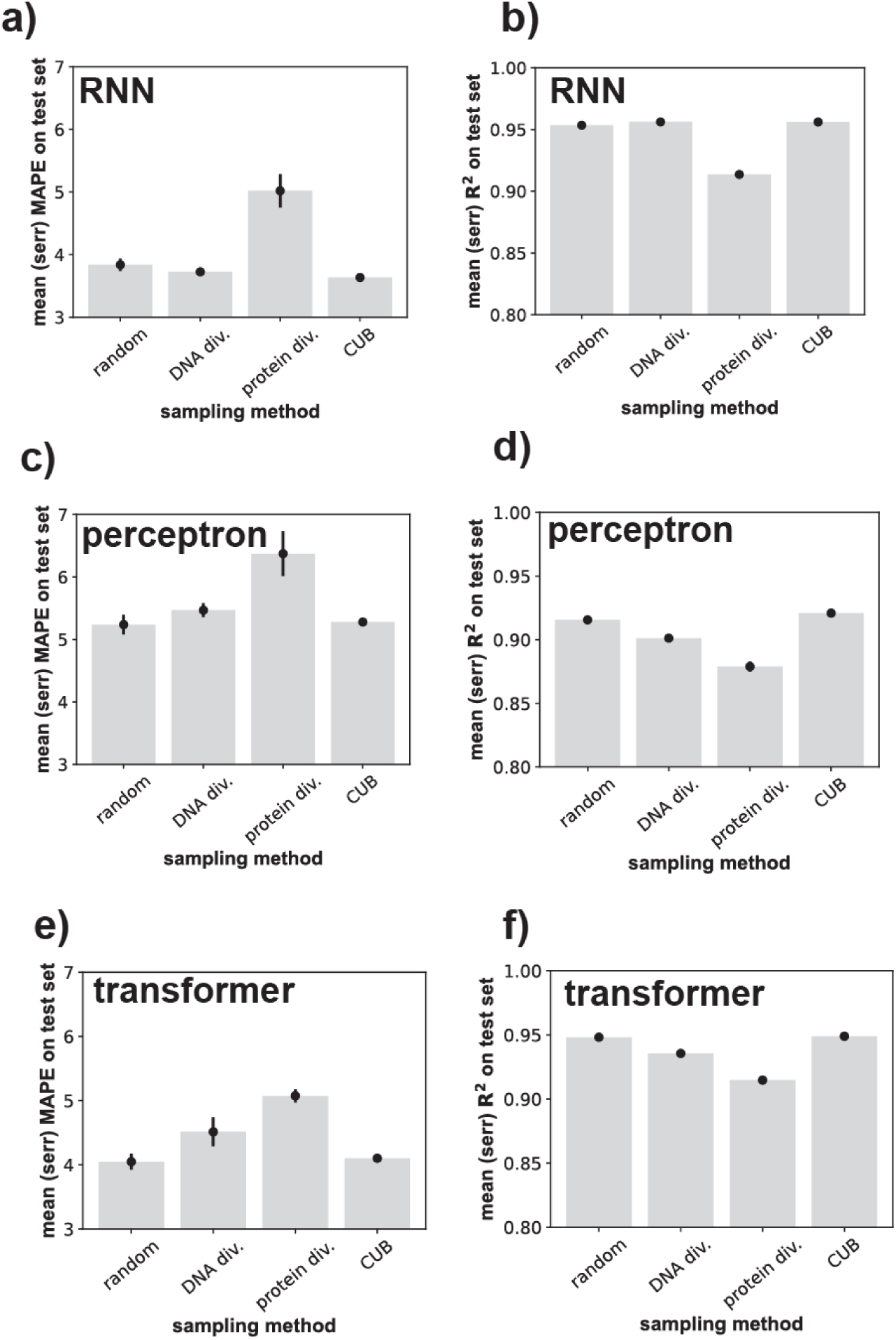
Sampling for amino acid sequence diversity substantially reduces RNN prediction quality for all architectures even at large sample sizes. Dot-whisker plots indicating the mean (height of bar) and standard error of the mean (whisker) of neural network prediction performance at a fixed sample size of *S*=8,000 genotypes for the four sampling methods shown on the horizontal axis. The *S* genotypes are either sampled randomly and uniformly (‘random’), to achieve maximal DNA sequence diversity (‘DNA div.’), maximal amino acid sequence diversity (‘protein dev.’), or maximal codon usage bias (‘CUB’, Methods). Panels **a)**, **c)**, and **e)** quantify prediction through the mape. Panels **b)**, **d)**, and **f)** quantify prediction through the coefficient of determination *R^2^*. a) and b) show prediction peerformance of the RNN; c) and d) show prediction performance of the multilayer perceptron; e) and f) show prediction performance of the transformer.

**Figure S11:**
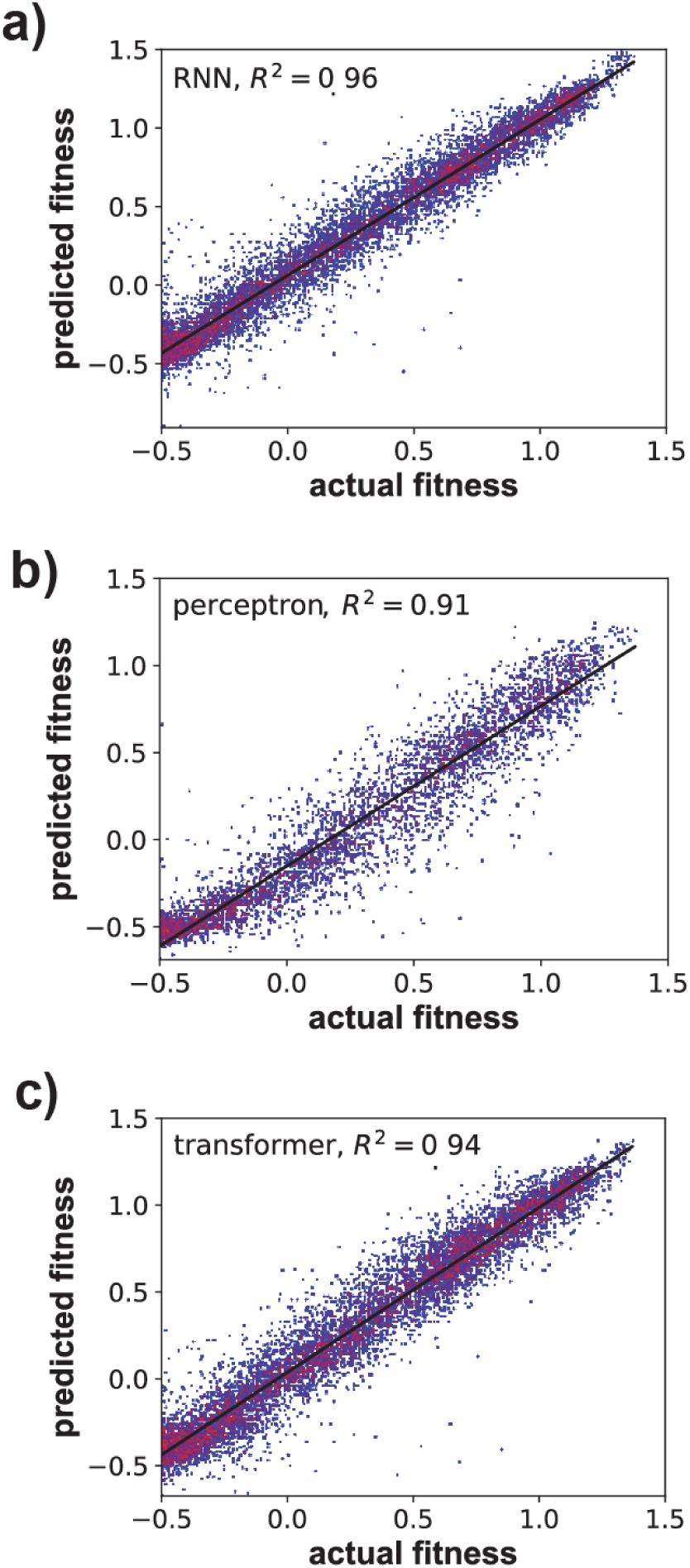
Measured and predicted fitness are highly correlated for the three best-performing deep learning architecture. I trained the best-performing (hypertuned) **a)** RNN, **b)** multilayer perceptron, and **c)** transformer architectures for 40 epochs on 25 percent of the viable genotypes, chosen randomly and with uniform probability among all 17,774 viable genotypes. The horizontal and vertical axis show measured and predicted fitness of viable genotypes, respectively, for a test data set comprising 50 percent of the remaining viable genotypes, also chosen randomly and uniformly. Colors indicate the density of data points from lowest (blue) to highest (red). The diagonal line is a linear regression line. A fitness value of zero corresponds to that of wild-type DHFR enzyme (low resistance to trimethoprim). The lowest viable fitness value equals -0.5 (Papkou, et al., 2023). Like elsewhere, I used the rmsprop algorithm for gradient descent during training, with minibatch sizes of 128 samples. Minor differences to values of *R^2^* reported in Table 1 are caused by stochastic differences in training that can arise during different training runs.

**Figure S12:**
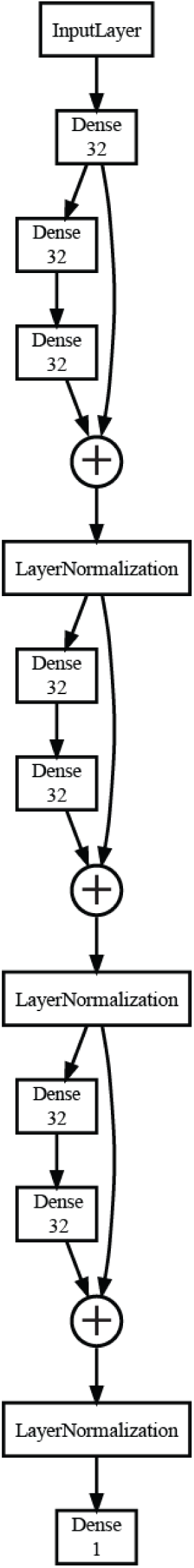
The best-performing multilayer perceptron architecture for regression. 7 dense layers of 32 output units each, followed by one dense output layer with a single unit, layer normalization after layer 3, 5, and 7, residual connection between output of layers 1 and 3, 3 and 5 (after layer normalization), as well as 5 and 7 (after layer normalization), no dropout, no regularization (see also Supplementary Results 1).

**Figure S13:**
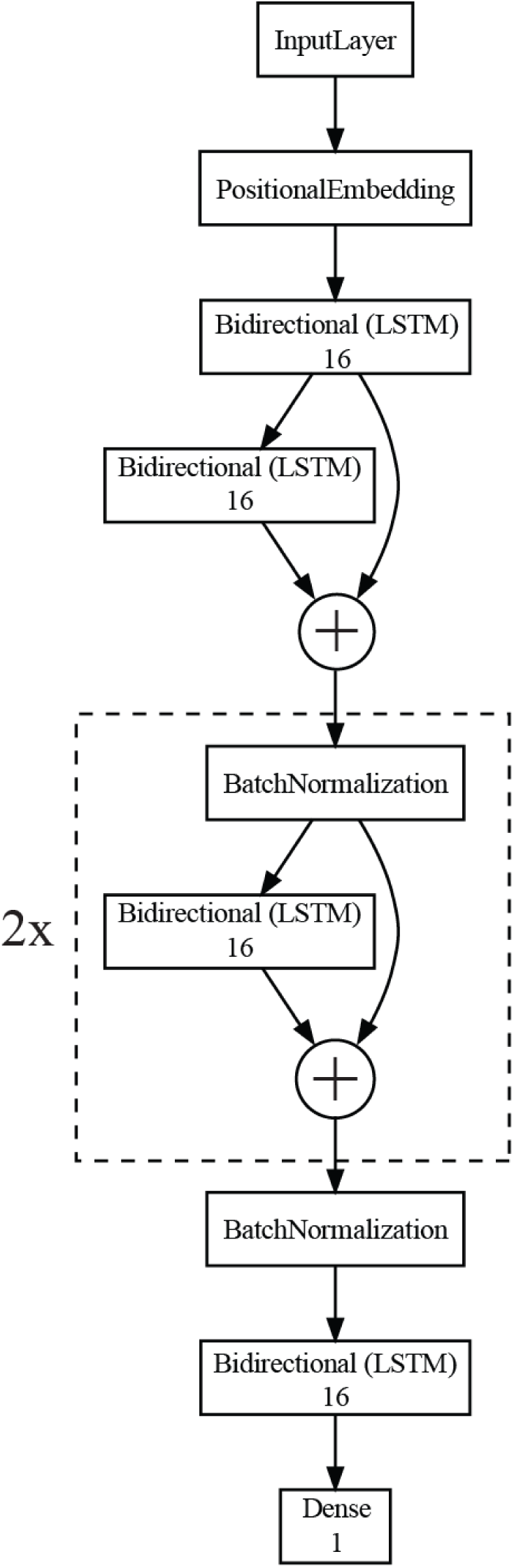
The best-performing RNN architecture for regression. The input data for each genotype consist of three integers, each representing an integer-encoded codon. These data are positionally embedded (Chollet, 2021, p 347) in a 32-dimensional embedding space. Each bidirectional LSTM recurrent neural networks has 16 recurrent units, uses a recurrent dropout of 0.1, but no recurrent or kernel regularization. ‘+’ indicates addition of outputs (residual connection). The last dense layer has a single unit (see also Supplementary Results 1).

**Figure S14:**
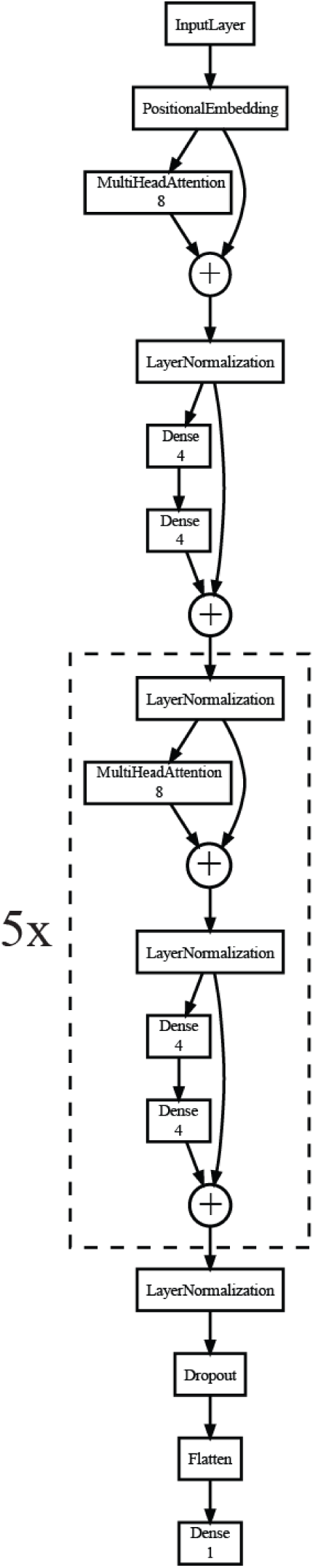
The best-performing transformer architecture for regression. The input data for each genotype consist of three integers, each representing an integer-encoded codon. These data are positionally embedded (Chollet, 2021, p 347) in a 32 dimensional embedding space, and form the input to the first of six identical transformer modules, each of which has eight attention heads, 16 key dimensions, and 16 units in the dense layer of each transformer module. A dropout of 0.1 is applied after the last transformer module, followed by one dense layer with a single unit. ‘+’ indicates addition of outputs (residual connection, see also Supplementary Results 1).

